# Allosteric Regulation of Pyruvate Kinase Enables Efficient and Robust Gluconeogenesis by Preventing Metabolic Conflicts and Carbon Overflow

**DOI:** 10.1101/2024.08.15.607825

**Authors:** Fukang She, Brent W. Anderson, Daven B. Khana, Shenwei Zhang, Wieland Steinchen, Danny K. Fung, Lauren N. Lucas, Nathalie G. Lesser, David M. Stevenson, Theresa J. Astmann, Gert Bange, Jan-Peter van Pijkeren, Daniel Amador-Noguez, Jue D. Wang

**Affiliations:** University of Wisconsin-Madison, Department of Bacteriology, Madison, USA; University of Wisconsin-Madison, Department of Food Science, Madison, USA; Philipps-University-Marburg, Center for Synthetic Microbiology (SYNMIKRO) & Faculty of Chemistry, Marburg, Germany

## Abstract

Glycolysis and gluconeogenesis are reciprocal metabolic pathways that utilize different carbon sources. Pyruvate kinase catalyzes the irreversible final step of glycolysis, yet the physiological function of its regulation is poorly understood. Through metabolomics and enzyme kinetics studies, we discovered that pyruvate kinase activity is inhibited during gluconeogenesis in the soil bacterium *Bacillus subtilis*. This regulation involves an extra C-terminal domain (ECTD) of pyruvate kinase, which is essential for autoinhibition and regulation by metabolic effectors. Introducing a pyruvate kinase mutant lacking the ECTD into *B. subtilis* resulted in defects specifically under gluconeogenic conditions, including inefficient carbon utilization, slower growth, and decreased resistance to the herbicide glyphosate. These defects are not caused by the phosphoenolpyruvate-pyruvate-oxaloacetate futile cycle. Instead, we identified two significant metabolic consequences of pyruvate kinase dysregulation during gluconeogenesis: increased carbon overflow into the medium and failure to expand glycolytic intermediates such as phosphoenolpyruvate (PEP). In silico analysis revealed that in wild-type cells, an expanded PEP pool enabled by pyruvate kinase regulation is critical for the thermodynamic feasibility of gluconeogenesis. Our findings underscore the importance of allosteric regulation during gluconeogenesis in coordinating metabolic flux, efficient energy utilization, and antimicrobial resistance.

## Introduction

Regulation of central carbon metabolism in microbes allows high efficiency usage of diverse carbon sources^1^. Central carbon metabolism involves glycolysis, gluconeogenesis, the pentose phosphate pathway, and the tricarboxylic acid (TCA) cycle. Glycolysis breaks glucose into two molecules of pyruvate, generating ATP and NADH and producing glycolytic intermediates essential for nucleotide and amino acid synthesis. In contrast, gluconeogenesis operates in the opposite direction to glycolysis and is required when carbon sources feeding into the early steps of glycolysis are not available. Gluconeogenesis produces essential glycolytic intermediates from non-sugar substrates including organic acids (e.g., malate, succinate, fumarate, pyruvate) and amino acids, by consuming energy generated from other pathways such as the TCA cycle.

Gluconeogenesis shares many enzymes with glycolysis for catalyzing reversible reactions, but it bypasses the irreversible reactions using different enzymes^2^. Shutting down irreversible glycolysis-specific reactions to avoid energy-wasting futile cycles is proposed to be crucial for gluconeogenesis^3^, however, experimental evidence remains scarce. The enzyme pyruvate kinase (EC 2.1.7.40) catalyzes the irreversible last step of glycolysis by transferring the phosphate from phosphoenolpyruvate (PEP) to ADP to generate ATP and pyruvate^4^. Pyruvate kinase has long been proposed to be inhibited during gluconeogenesis to avoid an energy-wasting futile cycle^5,6^, yet in bacteria it remains highly expressed during gluconeogenesis^7^. *In vitro*, pyruvate kinase is allosterically regulated by metabolic effectors such as fructose 1,6-bisphosphate (FBP) ^8,9^, AMP^9–14^, ribose 5-phosphate^9–12^ (R5P), glucose 6-phosphate (G6P) ^9,13,14^, and glycerol 3-phosphate (G3P) ^9^. Although allosteric regulations are believed to play a critical role in glycolysis, the physiological effects of pyruvate kinase regulation in bacteria are poorly understood due to a lack of dysregulated mutants.

In this study, we identified an essential autoinhibitory role of an extra C-terminal domain (ECTD) in pyruvate kinases, which is conserved across the bacterial phylum Firmicute (recently renamed Bacillota) ^12,15^. We demonstrated its crucial role in allosteric regulation of pyruvate kinase activity. Dysregulating pyruvate kinase with a domain-truncation mutant results in a constitutively active enzyme, enabling us to assess its physiological effects. Interestingly, this mutant exhibits no distinct phenotypes during glycolysis but manifests broad fitness defects specifically during gluconeogenesis. In contrast to the prevalent hypothesis of preventing an energy-wasting futile cycle, regulating pyruvate kinase activity prevents carbon overflow and increases PEP levels, promoting thermodynamically favorable and antimicrobial-resistant gluconeogenesis.

## Results

### The extra C-terminal domain is required for the inhibition of *B. subtilis* pyruvate kinase activity by metabolic effectors *in vitro*

Pyruvate kinase is a key glycolytic enzyme that shares conserved architecture of three principal domains (A, B, C domains) from bacteria to eukaryotes (Supplementary Fig. 1). In many bacterial species, pyruvate kinase also harbors an extra C-terminal domain (ECTD), which has been given various designations in previous studies, including ECTD, CT, ECTS, C′ domain, and PEPut^16–19^. Phylogenetic analysis of pyruvate kinase sequences from 113 NCBI reference genomes revealed that pyruvate kinases with an ECTD are conserved in the Bacillota phylum (except for *Streptococcus* species) and parts of Cyanobacteria phylum (Fig. 1a, Supplementary Table 1).

**Fig. 1:**
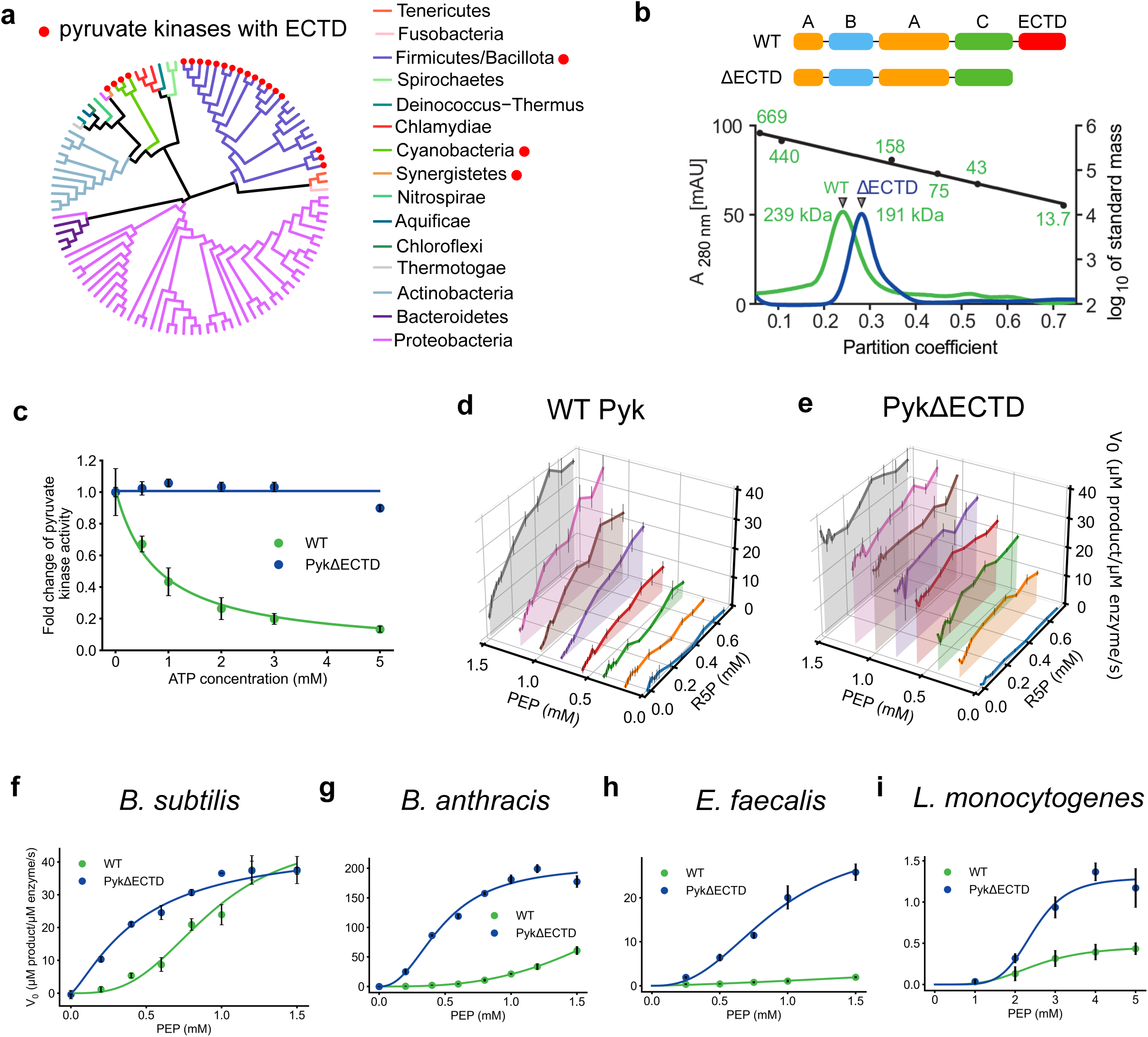
*In vitro* characterizations of enzymatic activities of full length pyruvate kinase (WT) and the ΔECTD variant (PykΔECTD). (a) Pyruvate kinases across a phylogenetic tree built from 113 bacterial reference genomes. Red dots indicate the presence of ECTD. Detailed species list is presented in Supplemental Table 1. (b) Size-exclusion chromatography of wild type (WT, green trace) and ΔECTD pyruvate kinase (blue trace) with apparent molecular masses of approximately 239 and 191 kDa, respectively. These correspond to the theoretical masses of 258.0 (monomeric mass of 64.5 kDa) and 212.8 kDa (monomeric mass of 53.2 kDa) for homoterameric wild type and ΔECTD pyruvate kinase. The black trace represents the average partition coefficients and linear regression thereof of a protein standard containing thyroglobulin (669 kDa), ferritin (440 kDa), aldolase (158 kDa), conalbumin (75 kDa), ovalbumin (43 kDa), and RNase A (13.7 kDa). (c) *B. subtilis* pyruvate kinase wild type the ΔECTD variant in the presence of indicated concentrations of ATP. Pyruvate kinase activity (the initial velocity of the reaction V_0_) is normalized against its activity in the absence of ATP. Error bars represent standard error of the mean of triplicates, unless otherwise stated. (d-e) Three dimensional plots of *B. subtilis* wild type (d) and ΔECTD (e) pyruvate kinase activities in the presence of combinations of different concentrations of the substrate PEP and activator R5P. (f-i) The *in vitro* enzymatic activities of wild type and ΔECTD pyruvate kinases from several representative Bacillota species, as indicated, at increasing concentrations of the substrate PEP (x-axis). Data were fitted to a nonessential activation equation (f) or an allosteric sigmoidal equation (g-i) to obtain the fitting curves (solid curves).

We investigated the impact of the pyruvate kinase ECTD using *Bacillus subtilis*, a well-established model bacterium within the Bacillota phylum. We recombinantly expressed and purified both the wild-type full length pyruvate kinase (WT Pyk) and a truncated variant lacking the ECTD domain (PykΔECTD) (Supplementary Fig. 2). Both the wild-type and truncated pyruvate kinase formed classical homotetramers in solution^6^, indicating that the ECTD is dispensable for pyruvate kinase oligomerization (Fig. 1b).

We measured pyruvate kinase activity using a classical coupled enzyme assay^20^. We first measured pyruvate kinase activity across various concentrations of its inhibitor, ATP^21^. Wild-type *B. subtilis* pyruvate kinase activity was significantly inhibited by ATP (<15% remaining at 5 mM ATP) (Fig. 1c). Given that ATP concentrations in *B. subtilis* cells typically reach around 3-5 mM under most growth conditions (e.g. Supplementary Table 3), all subsequent pyruvate kinase assays were conducted in the presence of 5 mM ATP.

Additionally, wild-type *B. subtilis* pyruvate kinase demonstrated activation by AMP or R5P (Supplemental Fig. 3a), known activators of previously characterized pyruvate kinases^11,12^. In contrast, the pyruvate kinase variant lacking ECTD maintained enzymatic activity but showed no response to either the inhibitor ATP (Fig. 1c), or the activators AMP and R5P (Supplemental Fig. 3b). This suggests that the ECTD domain is essential for regulating pyruvate kinase activity.

To systematically and quantitatively understand the regulation by metabolic effectors, we titrated both the substrate PEP and the activator R5P^12^ while keeping the inhibitor ATP at 5 mM. Pyruvate kinase activity was then measured for each combination of substrate and activator concentrations. Three-dimensional plots were generated with substrate concentration on the x-axis, activator concentration on the y-axis, and reaction rate on the z-axis (Fig. 1d, 1e). The plot revealed that wild-type pyruvate kinase exhibits a classical sigmoidal response to the substrate PEP^6^, with a Hill coefficient of 3.1, and is allosterically activated by R5P with a KA of 0.29 mM (Fig. 1d and Supplementary Fig. 3c, Table 1). In contrast, the pyruvate kinase variant lacking the ECTD domain shows activity even without the activator R5P and displays a Hill coefficient of 1.3 with respect to PEP (Fig. 1e and Supplementary Fig. 3d, Table 1).

**Table 1.**
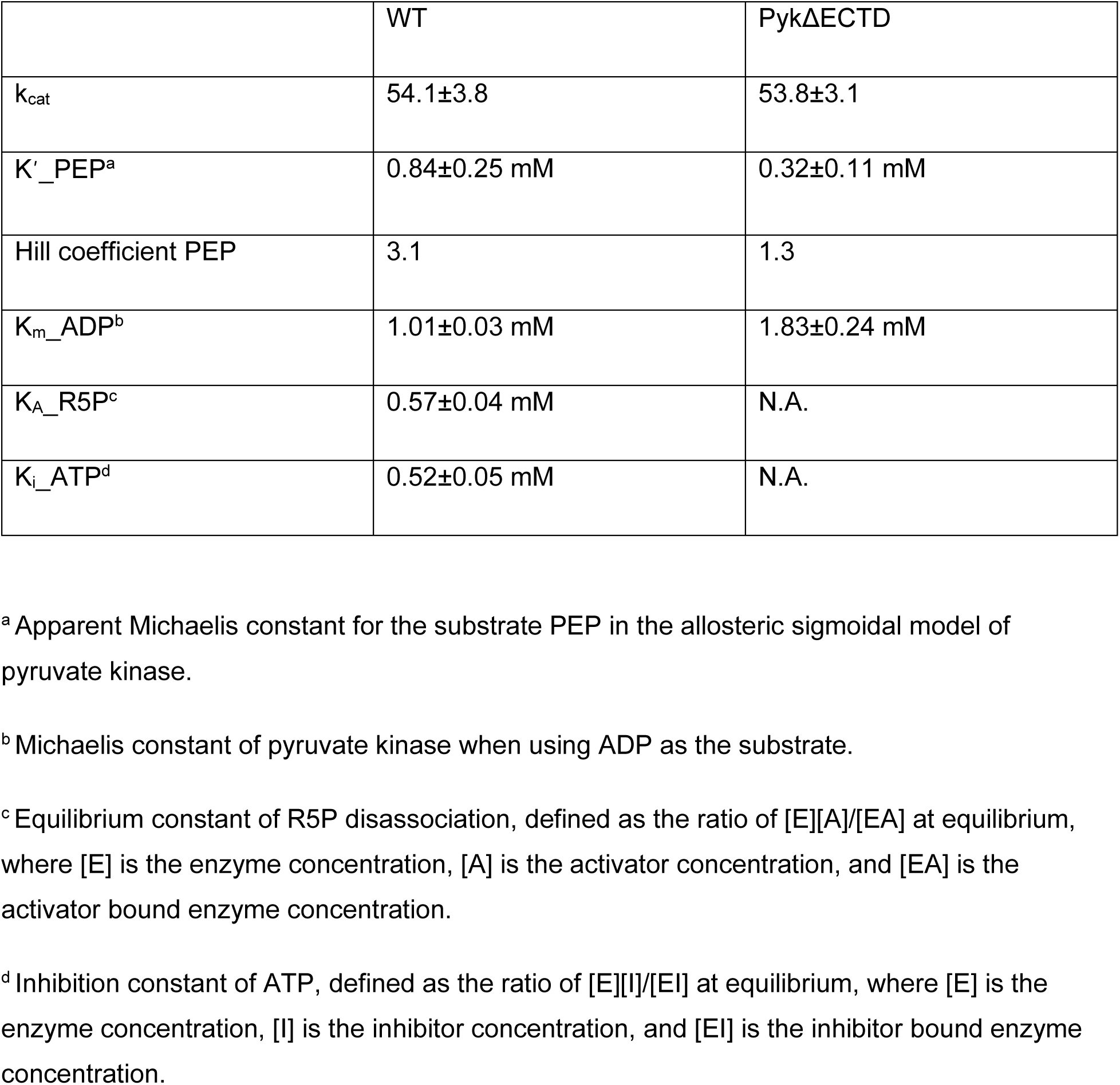
Enzyme kinetics parameters of *B. subtilis* pyruvate kinase.

**Table 2.**
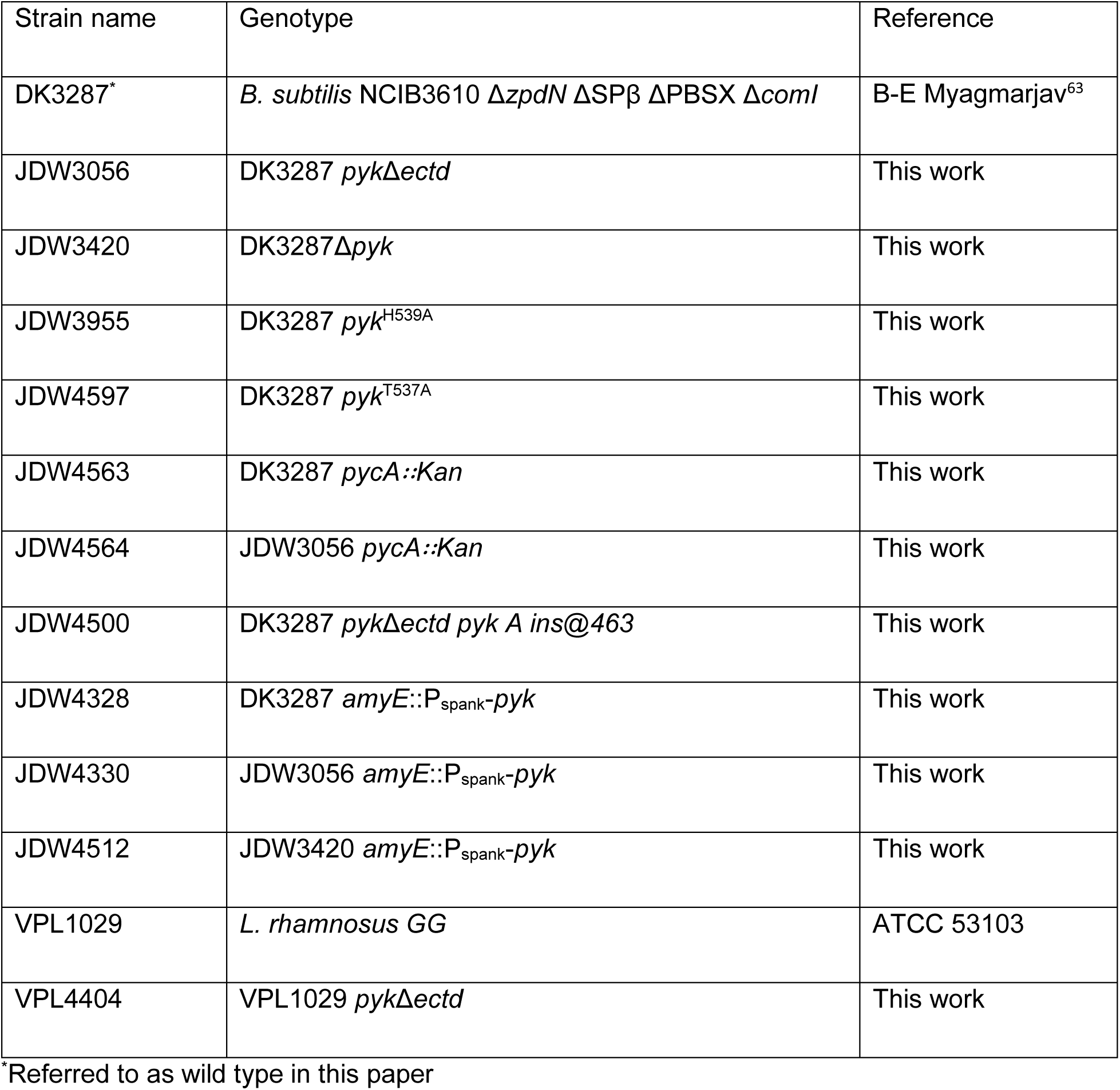
Strains used in this study.

**Table 3.** Plasmids used in this study.

| Plasmid name | Construct | Purpose | Reference |
| --- | --- | --- | --- |
| pJW269 | pLIC-trPC-HA | Expression vector | Stols L<sup>54</sup> |
| pJW422 | pLIC-trPC-HA-*pyk* | Expression of *B. subtilis* pyruvate kinase | This work |
| pJW724 | pLIC-trPC-HA-*pykΔectd* | Expression of *B. subtilis* ΔECTD pyruvate kinase (residues 1-474) | This work |
| pJW878 | pLIC-trPC-HA-Ban *pyk* | Expression of *B. anthracis* pyruvate kinase | This work |
| pJW879 | pLIC-trPC-HA-Ban *pykΔectd* | Expression of *B. anthracis* ΔECTD pyruvate kinase (residues 1-474) | This work |
| pJW880 | pLIC-trPC-HA-Efa *pyk* | Expression of *E. faecalis* pyruvate kinase | This work |
| pJW881 | pLIC-trPC-HA-Efa *pykΔectd* | Expression of *E. faecalis* ΔECTD pyruvate kinase (residues 1-474) | This work |
| pJW882 | pLIC-trPC-HA-Lmo *pyk* | Expression of *L. monocytogenes* pyruvate kinase | This work |
| pJW883 | pLIC-trPC-HA-Lmo *pykΔectd* | Expression of *L. monocytogenes* ΔECTD pyruvate kinase (residues 1-474) | This work |
| pJW557 | pPB41<sup>51</sup> without T7 promoter by the recombination template insertion site | CRISPR editing of genome | Anderson B<sup>64</sup> |
| pJW693 | pJW557 with *pykΔectd* guide RNA and repair template | CRISPR recombineering of *B. subtilis pykΔectd* | This work |
| pJW442 | pJW299<sup>65</sup> with 500 bp upstream and 500 bp downstream sequence of *B. subtilis pyk* | Engineering *B. subtilis Δpyk* | This work |
| pJW721 | pJW557 with *pyk*<sup>H539A</sup> guide RNA and repair template | CRISPR recombineering of *B. subtilis pyk*<sup>H539A</sup> | This work |
| pJW870 | pJW557 with *pyk*<sup>T537A</sup> guide RNA and repair template | CRISPR recombineering of *B. subtilis pyk*<sup>T539A</sup> | This work |
| pJW811 | pDR110 (from Rudner D) *amyE::P<sub>spank</sub>-B. subtilis pyk* | Express *B. subtilis* pyruvate kinase at non-endogenous site | This work |

To investigate whether the autoinhibitory role of the pyruvate kinase ECTD observed in *B. subtilis* is conserved across other Bacillota species, we purified pyruvate kinases and truncated ΔECTD variants from several representative species, including *Bacillus anthracis*, *Listeria monocytogenes*, and *Enterococcus faecalis*, and measured their activities. Although we observed that pyruvate kinases from different species showed varied responses to inhibitors and activators, the commonality (shown in Fig. 1f-i) is that wild type pyruvate kinases were consistently more inhibited compared to truncated ΔECTD variants within each species. These findings suggest that the ECTD domain acts as an autoinhibitory domain in representative Bacillota species, including *B. subtilis*, *B. anthracis, L. monocytogenes* and *E. faecalis*.

Although we observed that pyruvate kinase from different species showed varied responses to inhibitors and activators, the commonality (shown in Fig. 1f-i) is that wild-type pyruvate kinases were consistently more inhibited compared to truncated ΔECTD variants within each species.

### The extra C-terminal domain (ECTD) of pyruvate kinase is required for the inhibition of pyruvate kinase activity *in vivo*

Our enzymatic data show that pyruvate kinase ECTD is required for the regulation of pyruvate kinase activity by metabolic effectors *in vitro*. To evaluate the effect of ECTD in living bacterial cells, we engineered a *B. subtilis* mutant by replacing its wild-type pyruvate kinase gene (*pyk*) at its endogenous locus with a truncated *pyk* mutant retaining residues 1-474 but lacking ECTD (*pyk*Δ*ectd*). We then performed isotope tracing experiments using [1-^13^C] pyruvate as the sole carbon source. Cellular pyruvate can be derived either from external ^13^C-labeled pyruvate or from PEP via pyruvate kinase. Since PEP can be produced from both labeled oxaloacetate (OAA) from pyruvate and unlabeled OAA from the TCA cycle, increased pyruvate kinase activity would result in a higher concentration of unlabeled pyruvate in the cell.

By analyzing the isotope composition of primary metabolites in wild-type and *pyk*Δ*ectd* cells, we compared the *in vivo* activities of pyruvate kinase (Fig. 2a, Supplementary Fig. 4b). We found that the proportion of unlabeled PEP was similar in both wild type and mutant cells (Fig. 2b). However, the *pyk*Δ*ectd* mutant had a higher proportion of unlabeled pyruvate compared to wild-type cells (Fig. 2c), indicating higher pyruvate kinase activity in the *pyk*Δ*ectd* mutant. To rule out complications from pyruvate in the media being carried over during extraction, we also examined the isotope composition of alanine and valine, whose carbon backbones are derived entirely from pyruvate (Fig. 2d). Both alanine and valine in *pyk*Δ*ectd* cells had less ^13^C labeling than wild-type cells (Fig. 2e, 2f), confirming that pyruvate kinase activity is higher in the *pyk*Δ*ectd* mutant.

**Fig. 2:**
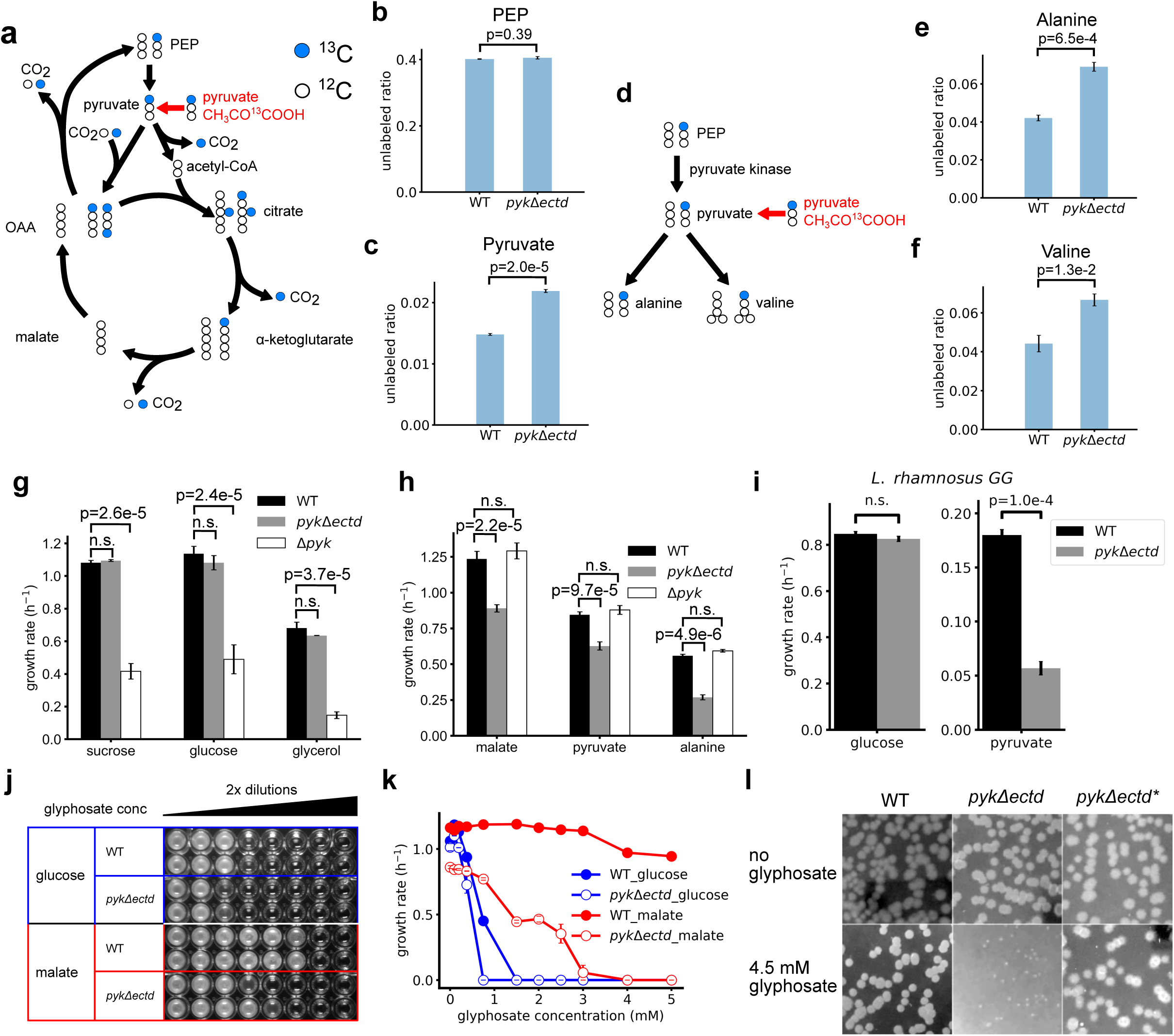
Metabolic flux, growth rate, and glyphosate resistance in full length (WT) and pyruvate kinase mutant lacking ECTD (*pyk*Δ*ectd*) during glycolysis and gluconeogenesis. (a) Schematics for quantification of pyruvate kinase activity *in vivo* by labeling major metabolites in cells grown with [1-^13^C] pyruvate. More unlabeled pyruvate and its derivatives will be observed if pyruvate kinase is more active. OAA: oxaloacetate. (b) Unlabeled ratio of PEP was the same in wild type cells and *pyk*Δ*ectd* cells. (c) Higher unlabeled ratio of pyruvate was observed in *pyk*Δ*ectd* cells compared to wild type cells grown in media with [1-^13^C] labeled pyruvate as the sole carbon source. (d) Carbon backbones of alanine and valine exclusively come from pyruvate, thus the ^13^C labeling of alanine and valine is a proxy of the intracellular pyruvate ^13^C labeling. (e-f) Higher unlabeled ratios of (f) alanine and (f) Valine were observed in *pyk*Δ*ectd* cells compared to wild type cells grown in media with [1-^13^C] labeled pyruvate as the sole carbon source. (g) Pyruvate kinase activity but not the ECTD is important for glycolytic growth. Steady state growth rates of *B. subtilis* with different alleles of pyruvate kinase, in different carbon sources (x-axis). WT: wild type cells with full length pyruvate kinase; *pyk*Δ*ectd*: pyruvate kinase mutant lacking ECTD; Δ*pyk*: deletion of pyruvate kinase. F test is performed for each group, followed by Tukey’s HSD test. p values of Tukey’s HSD test are labeled in the figure; p values larger than 0.05 are labeled as n.s. (not significant). (h) Pyruvate kinase activity is not required for gluconeogenic growth, but the ECTD is. (i) *L. rhamnosus GG pyk*Δ*ectd* mutant grew slower than the wild type strain in gluconeogenic media as well. (j) Minimal inhibitory concentration (MIC) of glyphosate for wild type and *pyk*Δ*ectd* cells in glycolytic or gluconeogenic media. *pyk*Δ*ectd* cells were more sensitive to glyphosate compared to the wild type cells, especially in gluconeogenic media. Glyphosate concentrations were 0.1875, 0.375, 0.75, 1.5, 3, 6, 12, 24 mM from left to right. (k) Growth rate of wild type and *pyk*Δ*ectd* strains in media with glucose or malate as the sole carbon source under different concentrations of glyphosate. Error bars represent standard error of the mean of duplicates. (l) A loss-of-function mutant of pyruvate kinase suppressed the hypersensitivity to glyphosate in *pyk*Δ*ectd* mutant.

### ECTD domain of pyruvate kinase promotes growth specifically during gluconeogenesis

Pyruvate kinase regulation by metabolic effectors has been proposed as a key mechanism for maintaining fitness during glycolysis and gluconeogenesis, but experimental evidence is lacking due to inability to engineer dysregulated pyruvate kinase mutants. The *pyk*Δ*ectd* mutant provides a crucial tool to evaluate the physiological consequences of failed pyruvate kinase inhibition. We first screened phenotypes of the *pyk*Δ*ectd* mutant on different carbon sources using Biolog Phenotype MicroArrays, which measure cell respiration as a reporter for growth (Supplementary Fig. 5). Intriguingly, the *pyk*Δ*ectd* mutant exhibited reduced total cell respiration rates on gluconeogenic carbon sources such as pyruvate and malate but not on glycolytic carbon sources such as glucose and fructose (Supplementary Fig. 5). The reduction in respiration can be explained if the *pyk*Δ*ectd* mutant population grew slower specifically during gluconeogenesis, resulting in fewer cells than wild type cells.

To test this hypothesis, we directly measured the growth curves of *B. subtilis* cells and compared the exponential growth rates of wild type pyruvate kinase (WT), pyruvate kinase deletion (Δ*pyk*), and the *pyk*Δ*ectd* mutant on either glycolytic (sucrose, glucose, glycerol) or gluconeogenic (pyruvate, malate, alanine) carbon sources (Fig. 2g, 2h). The Δ*pyk* mutant grew significantly slower than wild type cells during glycolytic growth (Fig. 2g), which is expected as pyruvate kinase activity is crucial for glycolysis. During growth on gluconeogenic carbon sources, the Δ*pyk* mutant grew similarly to wild type cells, confirming that pyruvate kinase activity is dispensable during gluconeogenesis (Fig. 2h).

In contrast to Δ*pyk,* there was no difference in growth rates between the wild-type and *pyk*Δ*ectd* strains during glycolysis, suggesting that pyruvate kinase regulation activity during glycolysis does not provide a fitness benefit. This somewhat contradicts the assumption that pyruvate kinase regulation is important for glycolysis. Strikingly, the *pyk*Δ*ectd* mutant grew significantly slower than wild type during gluconeogenesis (Fig. 2h), indicating that pyruvate kinase regulation is crucial for gluconeogenesis.

We performed complementation studies by overexpressing wild-type *pyk* from an ectopic locus, which successfully rescued the growth defect of the Δ*pyk* mutant during glycolysis (Supplementary Fig. 6a) but did not rescue the growth defect of the *pyk*Δ*ectd* mutant during gluconeogenesis (Supplementary Fig. 6b, 6c), confirming that *pyk*Δ*ectd* is a gain-of-function allele. Together with the isotope tracing experiments supporting ECTD-dependent regulation of pyruvate kinase activity during growth on the gluconeogenic carbon source pyruvate (Fig. 2a-h), these results suggest that the constitutively active pyruvate kinase leads to slow gluconeogenic growth.

To test whether pyruvate kinase ECTD plays a similar role in Bacillota beyond *B. subtilis*, we deleted the ECTD domain of pyruvate kinase in the probiotic Bacillota *Lacticaseibacillus rhamnosus GG*^22^. The resulting *L. rhamnosus GG pyk*Δ*ectd* mutant grew similarly to wild-type cells in glucose medium but displayed a growth defect in pyruvate medium (Fig. 2i, Supplemental Fig. 7), confirming that the pyruvate kinase ECTD domain also promotes gluconeogenic growth in *L. rhamnosus*.

### Bacteria display higher resistance to the herbicide glyphosate during gluconeogenesis than glycolysis dependent on pyruvate kinase ECTD

In addition to its impact on growth rate, we observed that the pyruvate kinase ECTD also enhances resistance to glyphosate, the active ingredient in the herbicide Roundup**^®^**^23^ and a potent antimicrobial compound^24^. Glyphosate competes with the glycolytic intermediate PEP to inhibit enolpyruvylshikimate 3-phosphate synthase and downstream aromatic amino acid synthesis^25,26^ (Supplementary Fig. 8a). *B. subtilis*, a soil bacterium, was sensitive to glyphosate when grown in glycolytic media, as reported previously^24^. However, to our surprise, *B. subtilis* displayed much higher resistance to glyphosate when the gluconeogenic malate was supplemented as the sole carbon source (Fig. 2j, 2k). Importantly, the protective effect of gluconeogenesis was strongly dependent on the ECTD domain of the pyruvate kinase. The minimal inhibitory concentration (MIC) of glyphosate for *pyk*Δ*ectd* cells was 4-fold lower than that for wild type cells (Fig. 2j) and the *pyk*Δ*ectd* mutant displayed strongly reduced growth rates compared to wild type cells (Fig. 2k). While wild type cells formed many colonies on solid media supplemented with 4.5 mM glyphosate and malate, *pyk*Δ*ectd* mutant cells formed no colonies except for a single suppressor (Fig. 2l, Supplementary Fig. 8b). Whole genome sequencing of this suppressor pinpointed a single mutation: a nucleotide frameshift insertion in the *pyk* locus at position 463 nt, causing a premature stop codon upstream of the active site, suggesting that loss of function of pyruvate kinase suppresses the sensitivity of *pyk*Δ*ectd* mutants to glyphosate. Thus, *B. subtilis* withstands glyphosate toxicity by enabling the negative regulation of pyruvate kinase activity during gluconeogenesis.

### Pyruvate kinase ECTD domain enhances carbon usage efficiency by preventing carbon overflow during gluconeogenesis

In addition to observing differences in growth rates, we also noticed differences in the final optical density between wild type and *pyk*Δ*ectd* cells. We cultured both strains in media containing limited amounts (2 g/L) of various carbon sources (glucose, malate, or pyruvate) and measured OD600 (optical density at 600 nm) as an indicator for cell biomass accumulation. When using glycolytic carbon sources, wild type and *pyk*Δ*ectd* cells reached similar maximal ODs (Fig. 3a, Supplementary Fig. 9a, 9b). However, with gluconeogenic carbon sources, wild type cells achieved 25% and 19% higher maximal OD compared to *pyk*Δ*ectd* cells in malate and pyruvate, respectively (Fig. 3a, Supplementary Fig. 9c, 9d). This suggests that *pyk*Δ*ectd* cells are less efficient in synthesizing biomass from gluconeogenic carbon sources compared to wild type cells.

**Fig 3.**
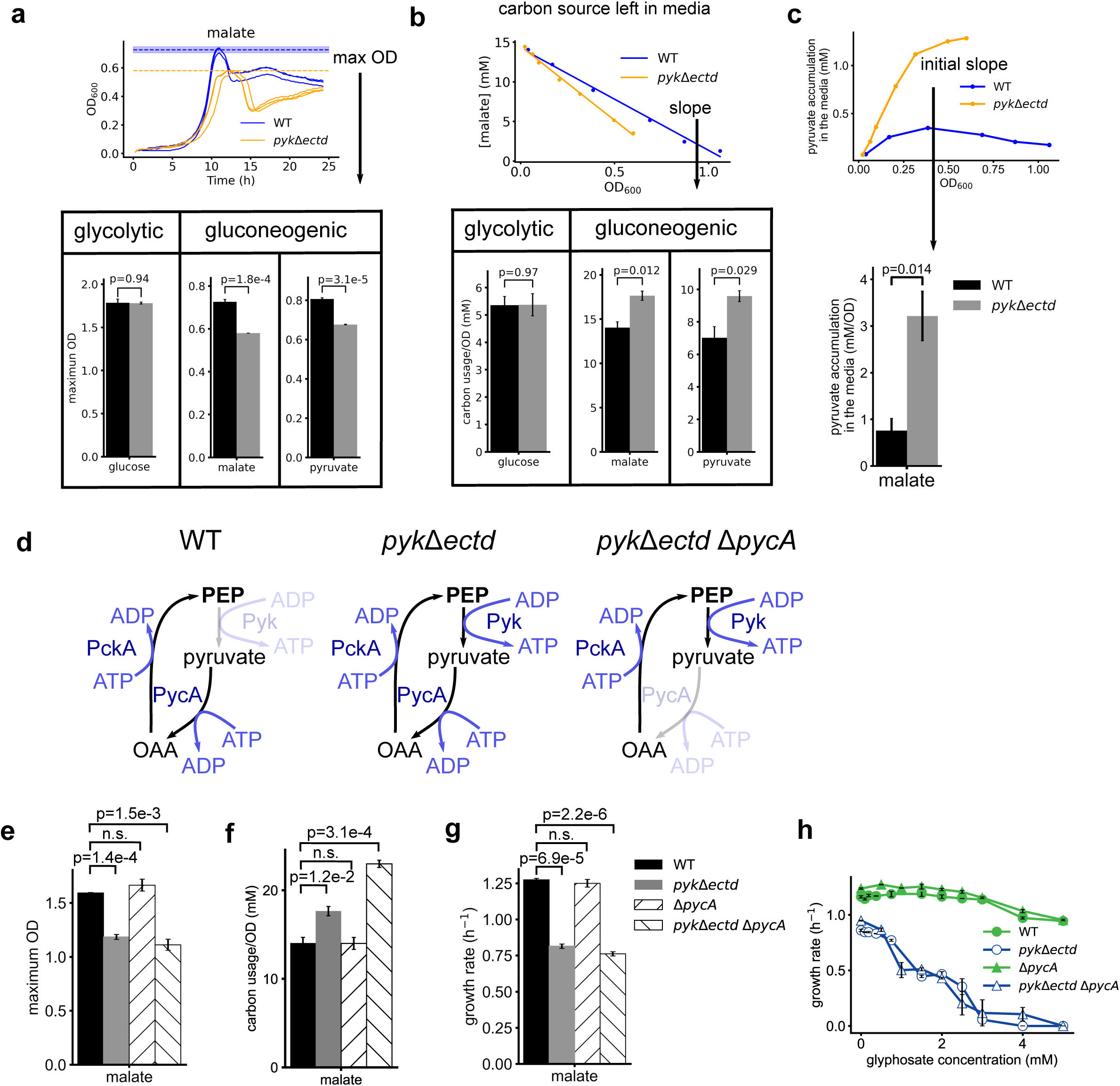
Loss of pyruvate kinase ECTD leads to low carbon use efficiency independent of the futile cycle during gluconeogenesis. (a) *pyk*Δ*ectd* cells had lower growth yield than the wild type cells. Maximum OD in media with limited carbon source was recorded as the proxy of biomass. (b) *pyk*Δ*ectd* cells consumed more gluconeogenic carbon sources than wild type cells per OD increase. Carbon source consumption rate was measured with HPLC or LC-MS. (c) *pyk*Δ*ectd* mutant excreted more pyruvate than the wild type strain when grew in media with malate as the sole carbon source. Pyruvate accumulation rate in the media was measured with LC-MS. (d) Schematics of the futile cycle model. Presumably, in *pyk*Δ*ectd* cells during gluconeogenic growth, pyruvate kinase, PycA and PckA are active simultaneously. The three reactions form a futile cycle that consumes one molecule of ATP every cycle without product accumulation. PycA: pyruvate carboxylase; PckA: PEP carboxykinase. (e-h) Disruption of the futile cycle does not rescue (e) and (f) the low gluconeogenic carbon use efficiency and (g) slow growth and (h) glyphosate hypersensitivity.

To directly measure carbon use efficiency, we quantified the consumption of carbon sources per unit of cell growth. We cultured wild type and *pyk*Δ*ectd* cells in media with glucose, malate, or pyruvate as the sole carbon source. During steady state growth, we used HPLC and LC-MS to measure the remaining carbon sources in the media (Supplementary Fig. 10). The results showed that *pyk*Δ*ectd* cells utilized glucose as efficiently as wild type cells but were significantly less efficient in utilizing malate and pyruvate (Fig. 3b). These findings confirm that *pyk*Δ*ectd* cells have lower carbon use efficiency, specifically during gluconeogenesis.

The reduced carbon use efficiency could be due to a futile carbon usage cycle, carbon overflow, or both. To determine if carbon overflow was occurring, we measured pyruvate excretion into the media using LC-MS when cells were grown with malate as the sole carbon source (Fig. 3c). We found that *pyk*Δ*ectd* cells excreted significantly more pyruvate and at a much higher rate (∼4X) compared to wild type cells. This overflow phenomenon likely contributes to the reduced efficiency of *pyk*Δ*ectd* cells in utilizing gluconeogenic carbon sources.

### Futile cycle does not contribute to the fitness defects of *pyk*Δ*ectd* during gluconeogenesis

A long-standing hypothesis is that pyruvate kinase is inhibited during gluconeogenesis to avoid an energy-wasting futile cycle^6^. In *B. subtilis*, pyruvate is converted to OAA by pyruvate carboxylase (PycA) and OAA is converted to PEP by PEP carboxykinase (PckA)^27^, with both enzymes active during gluconeogenesis. Failure to inhibit pyruvate kinase during gluconeogenesis would lead to PEP being converted back to pyruvate, forming a futile cycle that consumes ATP without product accumulation (Fig. 3d).

To test whether this futile cycle contributes to the gluconeogenic defects of *pyk*Δ*ectd*, we disrupted the cycle by deleting *pycA* (Fig. 3d), the gene encoding pyruvate carboxylase, in the *pyk*Δ*ectd* mutant. If the futile cycle among PEP, pyruvate and OAA contributes to the fitness defects, then removing *pycA* should rescue the defects observed in the *pyk*Δ*ectd* mutant. We assessed carbon utilization efficiency, growth rate, and glyphosate resistance in wild-type, *pyk*Δ*ectd*, Δ*pycA* and Δ*pycA pyk*Δ*ectd* double mutant in media with malate as the sole carbon source (Fig. 3e-h). Interestingly, Δ*pycA* did not rescue any of the defects introduced by *pyk*Δ*ectd* mutant. This indicates that the futile cycle does not contribute to the fitness defect of pyruvate kinase dysregulation during gluconeogenesis.

### Metabolomic analysis highlights the role of pyruvate kinase regulation in PEP pool expansion during gluconeogenesis

To understand the impact of pyruvate kinase regulation on central metabolism, we cultured wild type and mutant *B. subtilis* cells under both glycolytic and gluconeogenic conditions using glucose, malate, or pyruvate as the sole carbon source. We then monitored central carbon metabolites using LC-MS (Fig. 4, Supplementary Fig. 11, Supplementary Table 2). Wild-type cells grown in glucose exhibited significantly higher intracellular levels of upper glycolytic intermediates (e.g., G6P, FBP) and pentose phosphate pathway intermediates (e.g., R5P, S7P) compared to cells grown on malate or pyruvate. Conversely, levels of lower glycolytic intermediates (e.g., BPG, 3PG, PEP), TCA cycle intermediates (e.g., citrate), and amino acids derived from the TCA cycle (e.g., glutamate) were notably higher in cells grown on malate or pyruvate compared to glucose (Fig. 4b).

**Fig. 4:**
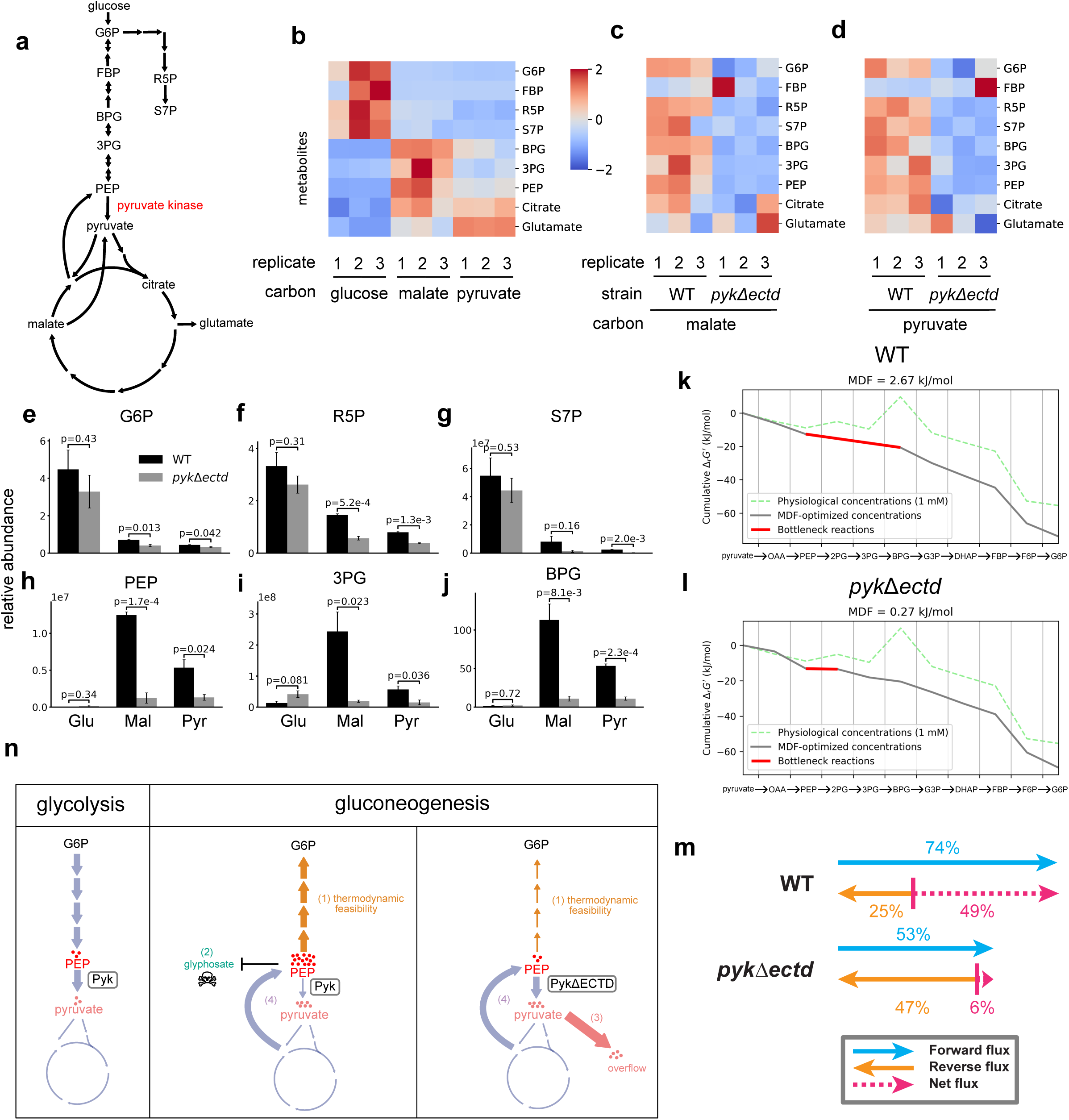
Loss of pyruvate kinase ECTD leads to PEP depletion and a less thermodynamically favorable gluconeogenesis. (a) Schematics of gluconeogenesis, part of the pentose phosphate pathway, and the TCA cycle. Important target metabolites are indicated. G6P: glucose 6-phosphate; FBP: fructose 1,6-bisphosphate; R5P: ribose 5-phosphate; S7P: sedoheptulose 7-phosphate; BPG: 1,3- bisphosphoglycerate; 3PG: 3-phosphoglycerate. (b) Z score of metabolites levels in wild type cells grown in glycolytic or gluconeogenic carbon sources. (c)-(d) Z score of metabolites levels in wild type and *pyk*Δ*ectd* cells grown in gluconeogenic carbon sources (c) malate and (d) pyruvate. (e)-(j) Relative abundance of metabolites upstream and downstream of pyruvate kinase. Relative abundances are calculated based on internal control, except for S7P, BPG and 3PG. See Materials and Methods: Metabolic analysis by LC-MS. (k)-(l) Cumulative drops in Gibbs free energies of gluconeogenesis for (k) wild type and (l) *pyk*Δ*ectd* cells. MDF: max-min driving force; F6P: fructose 6-phosphate; DHAP: dihydroxyacetone phosphate; G3P: glyceraldehyde 3-phosphate; 2PG: 2-phosphoglycerate. (m) The calculated forward flux ratio, reverse flux ratio and net flux ratio of the bottleneck reaction in gluconeogenesis for wild type and *pyk*Δ*ectd* cells grown in pyruvate. (n) A summary diagram of pyruvate kinase regulation and the physiological consequences of pyruvate kinase dysregulation during gluconeogenesis. (1): thermodynamic feasibility of gluconeogenesis; (2): glyphosate resistance; (3): pyruvate overflow; (4): the PEP-pyruvate-OAA futile cycle.

The metabolic profiles of wild type and *pyk*Δ*ectd* cells grown in glucose were comparable (Supplementary Fig. 11). However, major differences were observed in the *pyk*Δ*ectd* strain during growth on malate or pyruvate (Fig. 4c, 4d). Notably, PEP levels in the *pyk*Δ*ectd* mutant were significantly reduced, being more than three times lower than those in wild-type cells (Fig. 4h). Other lower-glycolytic intermediates (i.e., BPG, 3PG) were similarly depleted in the *pyk*Δ*ectd* mutant during gluconeogenesis, likely due to their equilibrium with PEP through reversible reactions (Fig. 4i, 4j). Additionally, G6P levels were reduced in *pyk*Δ*ectd* cells relative to wild-type cells during growth on malate and pyruvate (Fig. 4e). This decrease in G6P levels could be attributed to the role of PEP as an activator of the gluconeogenesis-specific enzyme fructose 1,6- bisphosphatase^28^.

Moreover, pentose phosphate pathway intermediates (R5P, S7P) were lower in *pyk*Δ*ectd* cells than in wild-type cells (Fig. 4f, 4g), likely due to the diminished levels of glycolytic intermediates that feed into this pathway. These findings suggests that the inability to deactivate pyruvate kinase in the *pyk*Δ*ectd* strain hampers PEP accumulation, negatively impacting gluconeogenic carbon flux and the generation of essential glycolytic intermediates for macromolecular biosynthesis.

### Expanded PEP pool is critical for the thermodynamic favorability of gluconeogenesis

To quantitatively determine if the reduced levels of PEP in the *pyk*Δ*ectd* mutant led to metabolic pathway bottlenecks, we integrated our metabolite concentration data with computational estimates of standard Gibbs free energies and performed a thermodynamic analysis of gluconeogenesis ^29^. To facilitate the modeling, we obtained the absolute intracellular metabolite concentrations using a previously described method^30^ in wild-type and *pyk*Δ*ectd* cells during growth on malate and pyruvate (Supplementary Table 3). We then used the Max-Min Driving Force (MDF) computational tool ^31^, which identifies the most thermodynamically constraining reaction(s) in a pathway and maximizes their reduction in Gibbs free energy using calculated optimized metabolite concentrations.

Our thermodynamic analyses revealed that in wild-type cells grown on pyruvate, the three reactions following PEP were identified as moderate thermodynamic bottlenecks, each with optimized free energies (MDF values) of -2.67 kJ/mol (Fig. 4k). However, in *pyk*Δ*ectd* cells grown on pyruvate, due to severely reduced levels of PEP, the reaction catalyzed by enolase (PEP to 2-phosphoglycerate) approached thermodynamic equilibrium (i.e., -0.27 kJ/mol) and became the single most constraining reaction step in gluconeogenesis (Fig. 4l).

To quantitate how this reduction in thermodynamic favorability impacts pathway flux, we used the flux-force efficacy relation (see Materials and Methods)^31^:

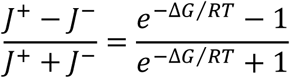

We determined that the decrease in free energy drop of the enolase reaction in *pyk*Δ*ectd* cells results in a severe nine-fold reduction in net flux compared to the net fluxes of the bottleneck reactions in wild-type cells (Fig. 4m). Consistent with these findings, enolase also became the single most thermodynamically constraining pathway step in gluconeogenesis in mutant cells grown on malate and exhibited a similar large reduction in net flux compared to wild-type cells (Supplementary Fig. 12).

Taken together, these results indicate that pyruvate kinase is inhibited during gluconeogenesis to help accumulate intracellular levels of PEP, making gluconeogenesis more thermodynamically favorable. However, pyruvate kinase lacking ECTD remains active, which leads to substantially lower intracellular levels of PEP, resulting in a more thermodynamically constrained gluconeogenesis and contributing to the observed slow growth.

## Discussion

In this study, we identified a regulatory mechanism that prevents metabolic conflict during gluconeogenesis (Fig. 4n). Specifically, pyruvate kinase, which converts PEP to pyruvate in glycolysis, is inhibited by its ECTD domain during gluconeogenesis, promoting efficient carbon utilization and faster growth rate by reducing carbon overflow and supporting PEP pool expansion. Conversely, mutants lacking this regulatory domain exhibit constitutively active pyruvate kinase, triggering simultaneous glycolytic and gluconeogenic reactions during gluconeogenesis, compromising carbon usage efficiency and slowing growth. Moreover, this dysregulation renders bacteria more vulnerable to glyphosate, which disrupts the synthesis of aromatic amino acids from PEP. The constitutive pyruvate kinase mutant fails to maintain high PEP levels, heightening its sensitivity to glyphosate. Thus, regulating pyruvate kinase is pivotal for metabolic coordination, significantly impacting carbon efficiency, growth optimization, and resistance to antimicrobial challenges.

Together these results reveal insights significantly different from what was known about pyruvate kinase regulation. In the literature, what is mostly known about allosteric regulation of pyruvate kinase has been focused on glycolysis. Instead, the understanding of pyruvate kinase regulation in gluconeogenesis exclusively revolves around its proposed role in preventing a futile cycle. However, our research demonstrates that preventing a futile cycle does not impact any of the three gluconeogenesis phenotypes we observed. Instead, we discovered that the thermodynamic uphill processes inherent in gluconeogenesis necessitate high PEP levels through pyruvate kinase regulation. These elevated PEP levels play a crucial role in facilitating timely generation of glycolytic intermediates, enabling resource allocation to important cellular processes such as nucleotide and cell envelope synthesis, while preventing inefficient overflow metabolism.

While it is clear how the constitutive pyruvate kinase mutant results in metabolic conflict, it remains unclear how ECTD enables the inhibition of enzyme activity during gluconeogenesis. Pyruvate kinases from many species have been shown to be allosterically regulated by multiple effectors^8–14,32^. The binding of allosteric effectors leads to the conformational change of the tetramer between inactive “T-state” and active “R-state”^14,33,34^. Pyruvate kinase ECTD interfaces with adjoining monomers in a crystal structure of tetrameric pyruvate kinase^15^. Based on our observations that ΔECTD pyruvate kinase remains as tetramers, it is most likely the ECTD is crucial for keeping the pyruvate kinase tetramer at “T-state” while allowing activation by effectors. In addition to mediating allosteric regulation, residues on ECTD may also undergo phosphorylation. ECTD shares homology with the swiveling domains (Pfam PF00391)^35^ of the pyruvate phosphate dikinase and several PEP-utilizing enzymes^36^ and contains a conserved active histidine residue that is phosphorylated by PEP or responsible for transferring phosphate group in other enzymes containing this domain^37–39^. This histidine (H539 in *B. subtilis*) is conserved in the Bacillota pyruvate kinase ECTD with the exception in *S. aureus* (Supplementary Fig. 1)^40^. Another potential phosphorylation site is threonine 537 on *B. subtilis* pyruvate kinase ECTD^41,42^. However, mutating either histidine 539 or threonine 537 to alanine, which inactivate potential phosphorylation of these sites, did not change the growth rate or yield regardless of growth conditions (Supplementary Fig. 13).

Although we show that ECTD serves an autoinhibitory function of pyruvate kinases from representative Bacillota species including *B. subtilis, B. anthracis, Listeria, E faecalis* and *L. rhamnosus*, there are exceptions. It is reported that ECTD promotes pyruvate kinase activity in *S. aureus*^14^ and does not have significant impact on pyruvate kinase activity from *Geobacillus thermophilus*^12,40^. However, the role of ECTD in these species has not been characterized *in vivo*. Furthermore, it should be note that many bacterial species beyond Bacillota phylum do not have ECTD, yet they are still allosterically regulated^8,10^. Finally, pyruvate kinase has been proposed to crosstalk with DNA replication enzymes^42^ and cell division in the Gram-positive bacterium *Bacillus subtilis*^43^. The multifaced impacts of pyruvate kinase in connection of the novel regulation we identified remain to be experimentally elucidated.

Our results do not support a hypothesis that pyruvate kinase is inhibited during gluconeogenesis to prevent an energy-wasting futile cycle^6^ among PEP, pyruvate and OAA. In a *B. subtilis* mutant with a constitutively active pyruvate kinase, we observed lower growth yield and slower growth rate during gluconeogenesis, the expected outcome from a futile cycle. However, these phenotypes persisted even when the futile cycle was disrupted, indicating that the futile cycle is not the cause of the defective gluconeogenesis upon pyruvate kinase dysregulation. Instead, the lower carbon use efficiency can be explained by an increased carbon overflow into the media; and the slow growth and glyphosate sensitivity can be explained by failure to build an expanded PEP pool. Similar result was found during glycolysis: while the formation of the PEP-pyruvate-OAA futile cycle was prevented by the CcpN-dependent repression of *pckA*^44^ and the constitutive mutant grew slower during glycolytic growth, the slow growth was the result of the depletion of the TCA cycle intermediates instead of the futile cycle^45^.

Together these results decoupled the importance of preventing a futile cycle from the role of pyruvate kinase regulation in building high levels of central carbon metabolic intermediate such as glycolytic and TCA cycle intermediates. We demonstrate that preventing a futile cycle does not significantly affect growth fitness. Instead, our research emphasizes the crucial role of maintaining high PEP levels to drive the reaction forward, ensuring an abundant supply of substrates. This enables efficient generation and strategic allocation of resources towards essential cellular processes, while minimizing diversion to TCA cycle and overflow metabolism.

Our results provide strong experimental validation for theoretical models that the inhibition of pyruvate kinase is important for anabolic metabolism^46^. In mammalian cells, it is proposed that constitutively active pyruvate kinase decreases the carbon flux channeling to biosynthetic pathways for nucleotides and some amino acids^47^. However, depletion of glycolytic intermediates is not directly observed in the mammalian metabolome during glycolytic growth^46^. Here we show with a bacterial model system, that a constitutively active pyruvate kinase does not affect the level of glycolytic intermediates during glycolytic growth but fails to build up high level of glycolytic intermediates during gluconeogenic growth, leading to defective anabolism. Thus, the role of pyruvate kinase regulation for proliferation, irrespective of details, may be conserved from bacteria to human.

Allosteric regulation at the enzymatic level is crucial for the survival of single-cell organisms in fluctuating nutrient conditions. *B. subtilis*, a soil bacterium, uses both glucose and malate as primary carbon sources, necessitating a rapid switch between gluconeogenesis and glycolysis for competitive fitness. By regulating pyruvate kinase activity, the bacterium can maintain high enzyme levels while swiftly adjusting to alternative carbon sources. Inhibiting pyruvate kinase through ECTD supports efficient gluconeogenesis and enables a quick response to changes in carbon availability. Future research in allosteric enzyme regulation will enhance our understanding of bacterial metabolic adaptation, with significant implications for bacterial growth and energy usage in natural environments.

## Materials and Methods

### Phylogenetic analysis

For the analysis of distribution of pyruvate kinases containing an ECTD, 118 bacterial reference genomes from the NCBI genome database were compared (Supplementary Table 1). Pyruvate kinase sequences were found in 113 out of 118 reference genomes based on annotations using Biopython^48^ and were aligned with MUSCLE in MEGA X^49^ using default settings. 16S rRNA sequences from these 113 genomes were identified based on the annotations using Biopython and aligned with ClustalW in MEGA X using default settings. A phylogenetic tree was built with MEGA X using the maximum likelihood method. Data were plotted with R package ggtree^50^.

### Plasmid and strain construction

A CRISPR/Cas9-based method^51^ was applied to engineer endogenous replacement of wild type *pyk* by *pyk*Δ*ectd* allele in *B. subtilis*. First, we modified the general purpose CRISPR/Cas9 vector pPB41^51^ by removing a T7 promoter upstream the repair template inserting site, preventing expression of the repair template sequence from expressing in *E. coli*, giving rise to pJW557. Next, the guide RNAs [oJW3089 (aaacACAGAAGAAGGCGGTTTGACTAGCCATGCTGg) and oJW3090 (aaaacCAGCATGGCTAGTCAAACCGCCTTCTTCTGT)], and the repair template (300 bp upstream and 300 bp downstream of ECTD) are inserted into pJW557 using Golden Gate Assembly (New England Biolabs), resulting in plasmid pJW724. Finally, pJW724 was transformed into wild type NCIB3610 strain (DK3287) and screened for *pyk*Δ*ectd* by Sanger sequencing of *pyk*, and by Illumina whole genome sequencing to verify that there were no second-site mutations.

For the deletion of pyruvate kinase, the plasmid pJW442 was constructed by inserting 500bp upstream and downstream of *pyk* into pEX44^52^. pJW442 was then transformed into DK3287, followed by transformation of I-sceI expression vector pJW296 to induce recombination, and colonies of the Δ*pyk* mutant were purified and confirmed by Sanger sequencing of PCR product.

VPL4404 (*L. rhamnosus GG pyk*Δ*ectd* retaining the N-terminal residues 1-477) was constructed by homologous recombination using dipeptide ligase as the counterselection marker^53^.

### Protein expression and purification

For pyruvate kinase expression and purification, full length pyruvate kinase or the ΔECTD variant (retaining the N-terminal residues 1-474 for *B. subtilis*, *B. anthracis, E. faecalis* or *L. monocytogenes* pyruvate kinase) coding sequences were cloned into the pLIC-trPC-HA vector^54^ (pJW269) downstream of a sequence encoding the hexa-histidine tag and the tobacco etch virus (TEV) protease cleavage site using ligation independent cloning (LIC)^54^. Expression vectors were transformed into *E. coli* BL21 (DE3) (NEB). A small-scale seed culture was grown in LB with 100μg/mL carbenicillin to OD600 around 0.8, and diluted 1:50 into fresh LB with 100μg/mL carbenicillin. 1 mM isopropyl β-D-1 thiogalactopyranoside (IPTG) (RPI) were added at around OD600 0.8 to induce the expression. After 4 hours of induction at 37°C, cells were pelleted and frozen at -80°C until purification.

Cells were suspended in lysis buffer (25 mM Tris-HCl pH=7.5, 300 mM NaCl, 10 mM imidazole) supplemented with 33 μg/mL deoxyribonuclease I to reduce the viscosity of the cell lysate, and lysed with a French press. The supernatant was collected after centrifugating the cell lysate, and filtered before loading to a HisTrap FF Column (Cytiva) on AKTA pure FPLC (Cytiva). The column was washed with 34.5 mM imidazole in lysis buffer for 5 column volumes and eluted with an increasing gradient of imidazole from 34.5 mM to 500 mM for 15 column volumes. His-tagged enzymes were digested with TEV protease to remove the hexa-histidine tag, and further purified by size-exclusion chromatography. Enzymes were concentrated to ∼8 mg/mL, and the enzyme concentration was measured by Bradford assay (Bio-Rad). Enzymes were stored in a buffer with 30 mM Tris-HCl (pH=7.5), 100 mM NaCl, 5 mM MgCl₂, 25 mM KCl, 1 mM DTT, and 10% (v/v) glycerol before being flash frozen with liquid nitrogen.

### Size exclusion chromatography

100 µl of 100 µM concentrated wild type or ΔECTD pyruvate kinase were applied onto a Superdex200 26/60 GL column (GE Healthcare) and eluted with SEC buffer (20 mM Tris-Cl pH 7.5, 300 mM NaCl, 1 mM EDTA, 1 mM DTT and 10% (v/v) glycerol) at 3 mL/min flow rate at 4 °C. A standard curve for molecular mass determination was obtained using a mixture of thyroglobulin (669 kDa), ferritin (440 kDa), aldolase (158 kDa), conalbumin (75 kDa), ovalbumin (43 kDa), and RNase A (13.7 kDa). The partition coefficient (Kav) was calculated as per Kav = (Ve-Vo)/(Vc-Vo) with the elution volumes (Ve) of the peak apex of the respective proteins, the given void volume of the column (Vo) of 100 ml and a total column volume (Vc) of 300 ml.

### Pyruvate kinase kinetics assay

For all *in vitro* assay from all species, pyruvate kinase activity was measured by an established coupled-enzyme assay with pyruvate kinase and lactate dehydrogenase^20^, in which the product of pyruvate kinase, pyruvate, is used by lactate dehydrogenase to convert NADH to NAD^+^. Production of NADH was monitored by the loss of NADH absorbance at 340 nm.

The *B. subtilis* pyruvate kinase reaction mix contained 100 mM Tris-HCl (pH=7.5), 100 mM KCl, 10 mM MgCl₂, 1.5 mM PEP (Millipore Sigma), 1 mM ADP (Millipore Sigma), 0.3 mM NADH (Millipore Sigma), and 25 U/mL L-lactate dehydrogenase (from bovine muscle, Millipore Sigma). When indicated, 1 mM AMP (Millipore Sigma) and/or 1 mM R5P (Biosynth International) and/or 5 mM ATP (Millipore Sigma) were added. Enzymatic reactions were initiated by addition of pyruvate kinase to a final concentration of 20 nM, and the absorbance at 340 nm was monitored in a quartz cuvette with a Shimadzu UV-2401PC spectrophotometer. Data was fitted using Python with Michaelis–Menten kinetics equations and plotted with Matplotlib^55^.

The *B. subtilis* pyruvate kinase activities in substrate and activator combinational titrations are conducted in 96 well plates and the absorbance at 340 nm was monitored by a Synergy 2 microplate reader (BioTek). The reaction mix contained 100 mM Tris-HCl (pH=7.5), 100 mM KCl, 10 mM MgCl₂, 0.5 mM ADP, 0.3 mM NADH, 25 U/mL L-lactate dehydrogenase, 5 mM ATP and different concentrations of PEP (0, 0.2, 0.4, 0.6, 0.8, 1.0, 1.2, 1.5 mM) and R5P (0, 20, 40, 60, 80, 100, 200, 300, 400, 500, 600, 700 µM) and 4 nM pyruvate kinase to start the reaction. Data were analyzed using Python and plotted with Matplotlib^55^. Wild type pyruvate kinase activity was normalized to the ΔECTD pyruvate kinase activity based on the maximum velocity. Both wild type and the ΔECTD pyruvate kinase kinetics data were fit to a nonessential activation equation that Hill coefficient was adapted to^56^:

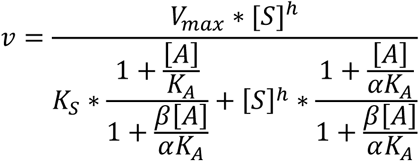

where [S] represents substrate concentration, [A] represents activator concentration, and h represents the Hill coefficient. Data were fit to the equation above using SciPy^57^.

Pyruvate kinases from other Bacillota species are tested with different concentrations of metabolic effectors for optimal activities. The *Bacillus anthracis* pyruvate kinase and ΔECTD variant reaction mix contained 1 mM AMP and 5 mM ATP. The *Enterococcus faecalis* pyruvate kinase and ΔECTD variant reaction mix contained 1 mM R5P. The *Listeria monocytogenes* pyruvate kinase and ΔECTD variant reaction mix contained 5 mM ATP.

### Biolog phenotypic screen

Wild type and *pyk*Δ*ectd* mutant were streaked on LB plates and grown overnight at 37°C. Cells were resuspended and diluted to an OD600 of 0.007 with inoculation fluid containing redox dye Dye Mix F (Biolog), and then 100 μL culture was inoculated into microplates PM1 and PM2 (Biolog) which contain different carbon sources. Absorbance at 590 nm was monitored by the OmniLog system for 24 hours at 37°C. Data was analyzed using Python and plotted with Matplotlib^55^.

### *B. subtilis* growth rate assay

To measure *B. subtilis* steady state growth rates, cells were streaked on LB plates and grown at 37°C overnight. 2-3 colonies were resuspended and inoculated into a modified S7 defined media^58^ [50 mM MOPS (pH adjusted to 7.0 with KOH), 10 mM (NH4)2SO4, 5 mM KH2PO4] (S750 minimal media), supplemented with 1% (w/v) carbon source [i.e., glucose, malic acid (pH adjusted to 7 with KOH), sodium pyruvate]. Cells were diluted and grown at 37°C overnight. Before reaching stationary phase, cells were sub-cultured into the same fresh media in 96-well plates with an initial OD600 ∼0.005, and OD600 was monitored by a Synergy 2 microplate reader (BioTek). Data was analyzed using Python and plotted with Matplotlib^55^.

For maximum growth density assays, cells were grown in the same way as for growth rate assays above, except that cells were washed with media without a carbon source before inoculation into media with 0.2% (w/v) carbon source to avoid carryover of nutrients.

### L. rhamnosus GG growth assay

*L. rhamnosus GG* wild type and *pyk*Δ*ectd* cells were inoculated in 5ml De Man, Rogasa and Sharpe (MRS, BD Difco) broth and incubated at 37°C for 16 hours. Overnight cultures were washed twice with modified MRS (10g peptone, 10g beef extract, 5g yeast extract, 2g ammonium citrate dibasic, 0.1g magnesium sulfate, 0.05g manganese sulfate, 2g dipotassium phosphate, and 1 mL Tween-80, dissolved in water to final volume 900 mL), followed by dilution into fresh mMRS supplemented with either 100 mM glucose, 100 mM of malate, or 100mM of pyruvate to OD600 = 0.05. Growth was monitored by a plate reader (Multiskan Sky, Thermo Fisher) at 37°C under hypoxic conditions (5% CO2, 2% O2).

### Glyphosate resistance assays

For glyphosate resistance assays, cells were grown in the same way as the growth rate assay, except that 0, 0.094, 0.188, 0.375, 0.75, 1.5,2, 2.5, 3, 4, 5 mM of glyphosate were added to the media. Glyphosate stock solution was made by dissolving glyphosate [N-(phosphonomethyl) glycine] (Sigma-Aldrich) in 100 mM KOH to 100 mM.

### Glyphosate resistance suppressors

JDW4500 was obtained by plating JDW3056 on an agar plate made from S750 minimal media supplemented with 1% (w/v) malic acid (pH adjusted to 7 with KOH), 1.5% (w/v) agar and 4.5 mM glyphosate. A single colony was obtained and verified. Genomic DNA was purified and sent for whole genome sequencing at SeqCenter. Mutation was identified from the sequencing result by breseq^59^.

### Carbon use efficiency measurement

To measure carbon usage efficiency, cells were grown in S750 minimal media supplemented with 0.2% (w/v) carbon source. As cells were growing, 2 mL of culture were centrifuged at 16100xg for 1 min to obtain supernatant for carbon source quantification, and 1 mL of culture was collected for OD measurement at each time point.

For quantification of glucose and pyruvate, media samples were mixed with 5 mM [U-¹³C] glucose or [1-¹³C] pyruvate as an internal control, and then quantified using LC-MS. MS parameters were set to a resolution of 140000, an automatic gain control (AGC) of 1e6, a maximum injection time of 40 ms, and a scan range of 70-1000 m/z. LC was performed on an ACQUITY UPLC BEH C18 column (1.7 μm, 2.1 × 100 mm; Waters). Total run time was 15 min with a flow rate of 0.2 mL/min, using 97:3 (v/v) water/methanol, 10 mM tributylamine (pH 8.2-8.5 adjusted with ∼9 mM acetic acid) as solvent A and 100% methanol as Solvent B. The gradient was as follows: 0 min, 5% B; 2.5 min, 5% B; 7.5 min, 95% B; 9.5 min, 95% B; 10 min, 5% B; 15 min, 5% B.

For quantification of malate, media samples were acidified with 18.4 mM H2SO4. After 10 min, samples were centrifuged at 16100 xg for 10 min, and the supernatant was loaded onto the HPLC. LC was performed on a Phenomenex Rezex ROA-Organic Acid H^+^ column. Total run time was 45 min with a flow rate of 0.3 mL/min, using 7.6 mM H2SO4 as the mobile phase. Refractive index was monitored for the quantification, and concentrations were calculated based on a standard curve generated with 0.8 mM, 4 mM and 20 mM malic acid standards.

Carbon source concentration in the media was plotted versus OD600, displaying a linear correlation. The slope, which was the carbon source consumed per unit of OD600, was used as the indicator of carbon use efficiency.

### Metabolomic analysis by LC-MS

Cells were grown in the same way as the growth rate assay. Cells from an overnight culture (exponential phase) were inoculated into the same fresh media (50 mM MOPS (pH adjusted to 7.0 with KOH), 10 mM (NH4)2SO4, 5 mM KH2PO4, and 1% (w/v) carbon source (i.e., glucose, malic acid (pH adjusted to 7 with KOH), sodium pyruvate)) in flasks at an initial OD600 ∼0.05 and grown to OD600 ∼0.5. 5 mL of culture were vacuum filtered to collect cells. Filter membranes with cells were submerged in 1.5 mL cold extraction solvent (acetonitrile: methanol: H₂O=40:40:20) over dry ice to quench metabolism. Cell extracts were centrifuged at 13200xg for 10 minutes to remove cell debris. The supernatant was then mixed with an internal control, dried by nitrogen gas flow, and resuspended in HPLC grade H₂O.

For preparation of the internal control, wild type cells were inoculated into media with 50 mM MOPS, 10 mM (NH4)2SO4, 5 mM KH2PO4, 1% (w/v) [U-¹³C] glucose, and grown at 37°C. Cells were inoculated into fresh [U-¹³C] glucose media in flasks at an initial OD600 ∼0.05 and grown to OD600 ∼0.5. Metabolites were extracted as described above.

For absolute metabolite concentration measurements, wild type or *pyk*Δ*ectd* cells were inoculated into media with 50 mM MOPS, 10 mM (NH4)2SO4, 5 mM KH2PO4, and 1% (w/v) [U-¹³C] glucose or 1% (w/v) [1-^13^C] pyruvate or natural malate, and grown in a 37°C shaker incubator overnight. Cells were inoculated into fresh corresponding media in flasks at an initial OD ∼0.05 and grown to OD ∼0.8. Unlabeled metabolites of interest were spiked in at 0x, 0.1x, 1x and 10x of estimated concentrations into the extraction solvent before extraction for glucose and pyruvate samples. For the malate samples, 0x spiked samples from the pyruvate group were mixed in at a 1:1 ratio. Cell extracts were dried by nitrogen gas flow, and resuspended in HPLC grade H2O.

To detect metabolites, samples were analyzed using HPLC-MS consisting of a Dionex UHPLC coupled by electrospray ionization (ESI, negative mode) to a Q Exactive Orbitrap mass spectrometer (Thermo Scientific) operated in full-scan mode for detection of targeted compounds based on their accurate masses. MS parameters were set to a resolution of 70000, an automatic gain control (AGC) of 1e6, a maximum injection time of 40 ms, and a scan range of 70-1000 m/z. LC was performed on an ACQUITY UPLC BEH C18 column (1.7 μm, 2.1 × 100 mm; Waters). Total run time was 25 min with a flow rate of 0.2 mL/min, using 97:3 (v/v) water/methanol, 10 mM tributylamine (pH 8.2-8.5 adjusted with ∼9 mM acetic acid) as solvent A and 100% methanol as solvent B. The gradient was as follows: 0 min, 5% B; 2.5 min, 5% B; 17 min, 95% B; 19.5 min, 95% B; 20 min, 5% B; 25 min, 5% B. Raw output data from MS were converted to mzXML format using in-house-developed software, and quantification of metabolites was performed using Metabolomic Analysis and Visualization Engine (MAVEN)^60,61^. Ion counts of metabolites were normalized to OD600 and internal controls when applicable. If the ion counts of a metabolite in the internal control was too low, then the ion counts of this metabolite would be normalized to OD600 only.

Z scores were calculated within different strains and carbon sources using the formula Z=(x-µ)/σ, where x is normalized ion counts, µ is the average of normalized ion counts of a certain metabolite within different conditions, and σ is the standard error of normalized ion counts of a certain metabolite within different conditions.

### Methods for Max-min driving force calculations

Optimized thermodynamic profiles for gluconeogenesis (2 Pyruvate è glucose 6-phosphate) were generated using the Max-min driving force (MDF) tool^31^ via the Python package equilibrator- pathway (version 0.5.0). Intracellular pH, ionic strength, and temperature were set to 7, 250 mM, and 298.15 K, respectively. Maximum and minimum concentration bounds for PEP, ATP, and ADP were based on a 40% range of the calculated absolute intracellular concentrations (Supplementary Table 3). For wild-type conditions, the concentration bounds for the remaining metabolites were based on default ranges of 1 μM-10 mM. To provide more conservative concentration ranges for the metabolites without absolute intracellular data for our *pyk*Δ*ectd* MDF profile, we combined the optimized metabolite concentrations generated from the wild-type model with our relative fold-change data for the gluconeogenic metabolites and set a 40% range on those metabolite concentrations. Forward and net flux ratios for wild-type and mutant conditions were calculated using the least thermodynamically favorable free energy of the respective pathways (i.e., the MDF value) and the following equation: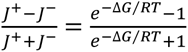^31,62^

## Acknowledgement

We thank Katrina Forest, Simon Gilroy, Srivatsan Raman, Atsushi Taguchi and members of the Wang and Amador-Noguez labs for discussions and comments on the manuscript. This work is supported by the USA National Institute of Health R35 GM127088 and Howard Hughes Medical Institute Faculty Scholar (to J.D.W.), the National Science Foundation (NSF) grant award no. 1715710 (to D.A.-N.), and the priority programme of the Deutsche Forschungsgemeinschaft (DFG) SPP1879 (to G.B.). S.Z. is a recipient of the Robert H. and Carol L. Deibel Distinguished Graduate Fellowship in Probiotic Research. B.W.A. was supported by NSF GRFP DGE-1256259.

## Author contributions

F.S., B.W.A., J.D.W. and D.A.N designed the research. F.S., S.Z., W.S., D.K.F., N.G.L., D.M.S. and T.J.A. performed experiments. F.S., D.B.K., S.Z., W.S., D.A.N and J.D.W analyzed data. F.S. wrote the paper. F.S., D.B.K., D.K.F, B.W.A, N.G.L, L.N.L, D.A.N and J.D.W edited the paper.

---

**Supplementary Fig. 1: Pyruvate kinases in some bacteria contain an extra C-terminal domain.**

Amino acid sequences alignment of pyruvate kinase in different species. Active site is in a pocket between A domain and B domain. C domain is a regulatory domain with effector binding sites. The active histidine on the extra C-terminal domain is circled by a rectangle, which is conserved in most Bacillota species but not *S. aureus*.

**Supplementary Fig. 2: Purification of *B. subtilis* pyruvate kinases.**

Coomassie brilliant blue stained SDS-PAGE gel of recombinantly expressed (a) wild type *Bacillus subtilis* pyruvate kinase and (b) ΔECTD *Bacillus subtilis* pyruvate kinase, purified with Ni-NTA column. L: lysate; W: wash.

**Supplementary Fig. 3: Three-dimensional analysis of pyruvate kinase kinetics: effects of substrate and activator concentrations on reaction rate.**

(a)-(b) Regulation of allosteric effectors on (a) wild type enzyme and (b) the ΔECTD enzyme variant. (c)-(d) Three-dimensional enzyme kinetic data in different combination of the substate PEP and activator R5P from Fig. 1d and 1e were fitted to an adapted nonessential activation equation^58^ by SciPy, and the fitted data were plotted with Matplotlib. Coefficient of determination (r^2^) was 0.98 for the fitting of wild type data and 0.97 for the fitting of PykΔECTD data. The catalytic rate constant and dissociation constants from the fit are listed in Table 1.

**Supplementary Fig. 4: Isotope composition of important metabolites in cells grown in media with [1-^13^C] pyruvate.**

(a) Schematics of the labeling experiment using [1-^13^C] pyruvate in Fig. 2. In *B. subtilis*, PckA, which converts OAA to PEP, is the only enzyme that regenerates PEP from TCA cycle intermediates. Therefore, when pyruvate is the sole carbon source, all PEP is produced from OAA. OAA can be synthesized from pyruvate directly or from malate. When cells are grown on [1-^13^C] pyruvate, OAA directly synthesized from pyruvate is [1-^13^C] labeled. However, OAA synthesized from malate after one round of the TCA cycle is unlabeled. Therefore, only a fraction of OAA is expected to be labeled, as well as PEP since it is derived from OAA. (b) Wild type and *pyk*Δ*ectd* cells were grown in media with [1-^13^C] pyruvate as sole carbon source, isotope composition of intracellular metabolites was analyzed by LC-MS. Isotope composition of some metabolites in gluconeogenesis and TCA cycle, and some amino acids are shown here. (c) Isotope composition of pyruvate in cells (left) and in the media (right).

**Supplementary Fig. 5: Phenotype profiling for *pyk*Δ*ectd* mutant.**

(a) Cells were inoculated in media with the indicated carbon sources and a redox dye which absorbs at 590 nm when reduced by cellular NADH as a reporter of growth. (b) Examples of several glycolytic (glucose, sucrose and fructose) and gluconeogenic carbon sources (malate, pyruvate and alanine) were plotted separately. Area under wild type curve is colored red, area under *pyk*Δ*ectd* curve is colored green, overlapping region is colored yellow.

**Supplementary Fig. 6: Complementation tests of *pyk* mutant alleles.**

Growth rates of wild type *pyk*, the *pyk*Δ*ectd* mutant, or the Δ*pyk* mutant alleles complemented with wild type *pyk* growing in glycolytic (Glucose, a) or gluconeogenic (malate, b or pyruvate, c) media. Wild type *pyk* was provided ectopically at the *amyE* locus with an IPTG-inducible promoter.

**Supplementary Fig. 7: Growth curve of *L. rhamnosus GG* in media with glycolytic or gluconeogenic carbon sources.**

Growth curves of the *L. rhamnosus GG* wild type and *pyk*Δ*ectd* cells in media with glucose or pyruvate as the sole carbon source. The same data as Fig. 2i.

**Supplementary Fig. 8: Hypersensitivity of *pyk*Δ*ectd* mutant to glyphosate can be suppressed by loss-of-function mutation of *pyk*.**

(a) Glyphosate inhibits the shikimate pathway which is required for the synthesis of aromatic amino acids by competing with PEP. EPSP: 5-enolpyruvylshikimate 3-phosphate; EPSPS: EPSP synthase. (b) Plate images of wild type, *pyk*Δ*ectd*, and glyphosate suppressor on malate minimal plates with or without glyphosate. The same data as Fig. 2l.

**Supplementary Fig. 9: Maximum OD600 of wild type or mutant cells grown in media with different sole carbon sources.**

Growth curve of wild type or *pyk*Δ*ectd* cells in media with 2 g/L (a) glucose, (b) glycerol, (c) malate or (d) pyruvate as the sole carbon source, dash lines represent the average maximum ODs for each condition, colored region represents 95% confidence interval.

**Supplementary Fig. 10: Gluconeogenic carbon use efficiency of *pyk*Δ*ectd* mutant is lower than wild type strain.**

Carbon sources remaining in spent media plotted versus OD600 of cells. Slopes are calculated by linear regression with the method of least squares, which represents carbon use efficiency.

**Supplementary Fig. 11: Metabolome of wild type and *pyk*Δ*ectd* mutant grown in glycolytic or gluconeogenic media.**

Heatmap of metabolites of wild type and *pyk*Δ*ectd* cells grown in media with different carbon sources. Rows and columns are clustered with UPGMA (unweighted pair group method with arithmetic mean). Color represents Z scores for rows. MEP: 2-C-methyl-D-erythritol 4- phosphate; GGPP: geranylgeranyl pyrophosphate.

**Supplementary Fig. 12: Thermodynamic analysis of gluconeogenesis for wild type and *pyk*Δ*ectd* cells grown in malate.**

(A)-(B) Cumulative drops in Gibbs free energies of gluconeogenesis for (a) wild type and (b) *pyk*Δ*ectd* cells. MDF: max-min driving force. (c) The calculated forward flux ration, reverse flux ratio and net flux ratio of the bottleneck reaction in gluconeogenesis for wild type and *pyk*Δ*ectd* cells grown in malate.

**Supplementary Fig. 13: Potential phosphorylation sites on pyruvate kinase ECTD are not responsible for defective gluconeogenic growth and low growth yield.**

(a) Growth rate of wild type, *pyk*Δ*ectd*, *pyk^H539A^* and *pyk^T537A^* cells in media with different sole carbon sources. (b) Maximum OD600 of wild type, *pyk*Δ*ectd*, *pyk^H539A^* and *pyk^T537A^* cells in media with malate as the sole carbon source.

## References

1. Buescher, J. M. et al. Global network reorganization during dynamic adaptations of Bacillus subtilis metabolism. Science 335, 1099–1103 (2012).

2. Cohen, G. N. Glycolysis, Gluconeogenesis and Glycogen Synthesis BT - Microbial Biochemistry: Second Edition. in (ed. Cohen, G. N.) 63–72 (Springer Netherlands, Dordrecht, 2011). doi:10.1007/978-90-481-9437-7_6.

3. Nelson 1942-, D. L. (David L. Lehninger Principles of Biochemistry. (Fourth edition. New York : W.H. Freeman, 2005., 2005).

4. Kayne, F. J. 11 Pyruvate Kinase. in Group Transfer Part A: Nucleotidyl Transfer Nucleosidyl Transfer Acyl Transfer Phosphoryl Transfer (ed. Boyer, P. D. B. T.-T. E.) vol. 8 353–382 (Academic Press, 1973).

5. Krebs, H. A. & Eggleston, L. V. THE ROLE OF PYRUVATE KINASE IN THE REGULATION OF GLUCONEOGENESIS. Biochem J 94, 3C–4C (1965).

6. Enriqueta Muñoz, Ma. & Ponce, E. Pyruvate kinase: current status of regulatory and functional properties. Comp Biochem Physiol B Biochem Mol Biol 135, 197–218 (2003).

7. Nicolas, P. et al. Condition-dependent transcriptome reveals high-level regulatory architecture in Bacillus subtilis. Science 335, 1103–1106 (2012).

8. Waygood, E. B. & Sanwal, B. D. The control of pyruvate kinases of Escherichia coli. I. Physicochemical and regulatory properties of the enzyme activated by fructose 1,6-diphosphate. J Biol Chem 249, 265–274 (1974).

9. Knowles, V. L., Smith, C. S., Smith, C. R. & Plaxton, W. C. Structural and Regulatory Properties of Pyruvate Kinase from the Cyanobacterium Synechococcus PCC 6301 *. Journal of Biological Chemistry 276, 20966–20972 (2001).

10. Waygood, E. B., Rayman, M. K. & Sanwal, B. D. The control of pyruvate kinases of Escherichia coli. II. Effectors and regulatory properties of the enzyme activated by ribose 5-phosphate. Can J Biochem 53, 444–454 (1975).

11. Sakai, H., Suzuki, K. & Imahori, K. Purification and properties of pyruvate kinase from Bacillus stearothermophilus. J Biochem 99, 1157–1167 (1986).

12. Zoraghi, R. et al. Functional Analysis, Overexpression, and Kinetic Characterization of Pyruvate Kinase from Methicillin-Resistant Staphylococcus aureus. Biochemistry 49, 7733–7747 (2010).

13. Noy, T. et al. Central Role of Pyruvate Kinase in Carbon Co-catabolism of Mycobacterium tuberculosis. J Biol Chem 291, 7060–7069 (2016).

14. Zhong, W. et al. Allosteric pyruvate kinase-based “logic gate” synergistically senses energy and sugar levels in Mycobacterium tuberculosis. Nat Commun 8, 1986 (2017).

15. Suzuki, K., Ito, S., Shimizu-Ibuka, A. & Sakai, H. Crystal structure of pyruvate kinase from Geobacillus stearothermophilus. J Biochem 144, 305–312 (2008).

16. Enriqueta Muñoz, Ma. & Ponce, E. Pyruvate kinase: current status of regulatory and functional properties. Comp Biochem Physiol B Biochem Mol Biol 135, 197–218 (2003).

17. Nguyen, C. C. & Saier, M. H. Phylogenetic analysis of the putative phosphorylation domain in the pyruvate kinase of Bacillus stearothermophilus. Res Microbiol 146, 713–719 (1995).

18. Suzuki, K., Ito, S., Shimizu-Ibuka, A. & Sakai, H. Crystal structure of pyruvate kinase from Geobacillus stearothermophilus. J Biochem 144, 305–312 (2008).

19. Horemans, S. et al. Pyruvate kinase, a metabolic sensor powering glycolysis, drives the metabolic control of DNA replication. BMC Biol 20, 1–26 (2022).

20. Kornberg, A. & Pricer, W. E. ENZYMATIC PHOSPHORYLATION OF ADENOSINE AND 2,6-DIAMINOPURINE RIBOSIDE. Journal of Biological Chemistry 193, 481–495 (1951).

21. Sakai, H., Suzuki, K. & Imahori, K. Purification and properties of pyruvate kinase from Bacillus stearothermophilus. J Biochem 99, 1157–1167 (1986).

22. Doron, S., Snydman, D. R. & Gorbach, S. L. Lactobacillus GG: bacteriology and clinical applications. Gastroenterol Clin North Am 34, 483–98, ix (2005).

23. Duke, S. O. & Powles, S. B. Glyphosate: a once-in-a-century herbicide. Pest Manag Sci 64, 319– 325 (2008).

24. Fischer, R. S., Berry, A., Gaines, C. G. & Jensen, R. A. Comparative action of glyphosate as a trigger of energy drain in eubacteria. J Bacteriol 168, 1147–1154 (1986).

25. Fischer, R. S., Rubin, J. L., Gaines, C. G. & Jensen, R. A. Glyphosate sensitivity of 5-enol- pyruvylshikimate-3-phosphate synthase from Bacillus subtilis depends upon state of activation induced by monovalent cations. Arch Biochem Biophys 256, 325–334 (1987).

26. Steinrücken, H. C. & Amrhein, N. The herbicide glyphosate is a potent inhibitor of 5-enolpyruvyl- shikimic acid-3-phosphate synthase. Biochem Biophys Res Commun 94, 1207–1212 (1980).

27. Sauer, U. & Eikmanns, B. J. The PEP-pyruvate-oxaloacetate node as the switch point for carbon flux distribution in bacteria. FEMS Microbiol Rev 29, 765–794 (2005).

28. Fujita, Y. & Freese, E. Purification and properties of fructose-1,6-bisphosphatase of Bacillus subtilis. J Biol Chem 254, 5340–5349 (1979).

29. Noor, E., Haraldsdóttir, H. S., Milo, R. & Fleming, R. M. T. Consistent estimation of Gibbs energy using component contributions. PLoS Comput Biol 9, e1003098 (2013).

30. Bennett, B. D., Yuan, J., Kimball, E. H. & Rabinowitz, J. D. Absolute quantitation of intracellular metabolite concentrations by an isotope ratio-based approach. (2008) doi:10.1038/nprot.2008.107.

31. Noor, E. et al. Pathway thermodynamics highlights kinetic obstacles in central metabolism. PLoS Comput Biol 10, e1003483 (2014).

32. Mattevi, A. et al. Crystal structure of Escherichia coli pyruvate kinase type I: molecular basis of the allosteric transition. Structure 3, 729–741 (1995).

33. Morgan, H. P. et al. Allosteric mechanism of pyruvate kinase from Leishmania mexicana uses a rock and lock model. J Biol Chem 285, 12892–12898 (2010).

34. Morgan, H. P. et al. M2 pyruvate kinase provides a mechanism for nutrient sensing and regulation of cell proliferation. Proc Natl Acad Sci U S A 110, 5881–5886 (2013).

35. Nguyen, C. C. & Saier, M. H. Phylogenetic analysis of the putative phosphorylation domain in the pyruvate kinase of Bacillus stearothermophilus. Res Microbiol 146, 713–719 (1995).

36. Pocalyko, D. J., Carroll, L. J., Martin, B. M., Babbitt, P. C. & Dunaway-Mariano, D. Analysis of sequence homologies in plant and bacterial pyruvate phosphate dikinase, enzyme I of the bacterial phosphoenolpyruvate: sugar phosphotransferase system and other PEP-utilizing enzymes. Identification of potential catalytic and regulatory motif. Biochemistry 29, 10757– 10765 (1990).

37. Lim, K. et al. Swiveling domain mechanism in pyruvate phosphate dikinase. Biochemistry 46, 14845–14853 (2007).

38. Teplyakov, A. et al. Structure of phosphorylated enzyme I, the phosphoenolpyruvate:sugar phosphotransferase system sugar translocation signal protein. Proc Natl Acad Sci U S A 103, 16218–16223 (2006).

39. Herzberg, O. et al. Swiveling-domain mechanism for enzymatic phosphotransfer between remote reaction sites. Proc Natl Acad Sci U S A 93, 2652–2657 (1996).

40. Sakai, H. Possible structure and function of the extra C-terminal sequence of pyruvate kinase from Bacillus stearothermophilus. J Biochem 136, 471–476 (2004).

41. Eymann, C. et al. Dynamics of protein phosphorylation on Ser/Thr/Tyr in Bacillus subtilis. Proteomics 7, 3509–3526 (2007).

42. Horemans, S. et al. Pyruvate kinase, a metabolic sensor powering glycolysis, drives the metabolic control of DNA replication. BMC Biol 20, 1–26 (2022).

43. Monahan, L. G., Hajduk, I. V, Blaber, S. P., Charles, I. G. & Harry, E. J. Coordinating bacterial cell division with nutrient availability: a role for glycolysis. mBio 5, e00935–14 (2014).

44. Servant, P., Le Coq, D. & Aymerich, S. CcpN (YqzB), a novel regulator for CcpA-independent catabolite repression of Bacillus subtilis gluconeogenic genes. Mol Microbiol 55, 1435–1451 (2005).

45. Tännler, S. et al. CcpN controls central carbon fluxes in Bacillus subtilis. J Bacteriol 190, 6178– 6187 (2008).

46. Israelsen, W. J. & Vander Heiden, M. G. Pyruvate kinase: Function, regulation and role in cancer. Semin Cell Dev Biol 43, 43–51 (2015).

47. Lunt, S. Y. et al. Pyruvate kinase isoform expression alters nucleotide synthesis to impact cell proliferation. Mol Cell 57, 95–107 (2015).

48. Cock, P. J. A. et al. Biopython: freely available Python tools for computational molecular biology and bioinformatics. Bioinformatics 25, 1422–1423 (2009).

49. Kumar, S., Stecher, G., Li, M., Knyaz, C. & Tamura, K. MEGA X: Molecular Evolutionary Genetics Analysis across Computing Platforms. Mol Biol Evol 35, 1547–1549 (2018).

50. Yu, G., Smith, D. K., Zhu, H., Guan, Y. & Lam, T. T.-Y. ggtree: an r package for visualization and annotation of phylogenetic trees with their covariates and other associated data. Methods Ecol Evol 8, 28–36 (2017).

51. Burby, P. E. & Simmons, L. A. CRISPR/Cas9 Editing of the Bacillus subtilis Genome. Bio Protoc 7, e2272 (2017).

52. Comella, N. & Grossman, A. D. Conservation of genes and processes controlled by the quorum response in bacteria: characterization of genes controlled by the quorum-sensing transcription factor ComA in Bacillus subtilis. Mol Microbiol 57, 1159–1174 (2005).

53. Zhang, S., Oh, J.-H., Alexander, L. M., Özçam, M. & van Pijkeren, J.-P. d-Alanyl-d-Alanine Ligase as a Broad-Host-Range Counterselection Marker in Vancomycin-Resistant Lactic Acid Bacteria. J Bacteriol 200, (2018).

54. Stols, L. et al. A new vector for high-throughput, ligation-independent cloning encoding a tobacco etch virus protease cleavage site. Protein Expr Purif 25, 8–15 (2002).

55. Hunter, J. D. Matplotlib: A 2D Graphics Environment. Comput Sci Eng 9, 90–95 (2007).

56. Segel, I. H. Enzyme Kinetics : Behavior and Analysis of Rapid Equilibrium and Steady State Enzyme Systems. (Wiley, New York, 1993).

57. Virtanen, P. et al. SciPy 1.0: fundamental algorithms for scientific computing in Python. Nat Methods 17, 261–272 (2020).

58. Vasantha, N. & Freese, E. Enzyme changes during Bacillus subtilis sporulation caused by deprivation of guanine nucleotides. J Bacteriol 144, 1119–1125 (1980).

59. Deatherage, D. E. & Barrick, J. E. Identification of mutations in laboratory-evolved microbes from next-generation sequencing data using breseq. Methods Mol Biol 1151, 165–188 (2014).

60. Clasquin, M. F., Melamud, E. & Rabinowitz, J. D. LC-MS data processing with MAVEN: a metabolomic analysis and visualization engine. Curr Protoc Bioinformatics **Chapter** 14, Unit14.11 (2012).

61. Melamud, E., Vastag, L. & Rabinowitz, J. D. Metabolomic analysis and visualization engine for LC- MS data. Anal Chem 82, 9818–9826 (2010).

62. Khana, D. B., Callaghan, M. M. & Amador-Noguez, D. Novel computational and experimental approaches for investigating the thermodynamics of metabolic networks. Curr Opin Microbiol 66, 21–31 (2022).

63. Myagmarjav, B.-E., Konkol, M. A., Ramsey, J., Mukhopadhyay, S. & Kearns, D. B. ZpdN, a Plasmid- Encoded Sigma Factor Homolog, Induces pBS32-Dependent Cell Death in Bacillus subtilis. J Bacteriol 198, 2975–2984 (2016).

64. Anderson, B. W. et al. The nucleotide messenger (p)ppGpp is an anti-inducer of the purine synthesis transcription regulator PurR in Bacillus. Nucleic Acids Res 50, 847–866 (2022).

65. Liu, K. et al. Molecular mechanism and evolution of guanylate kinase regulation by (p)ppGpp. Mol Cell 57, 735–749 (2015).

