## Supplementary Figures and Tables for "Allosteric Regulation of Pyruvate Kinase Enables Efficient and Robust Gluconeogenesis by Preventing Metabolic Conflicts and Carbon Overflow"

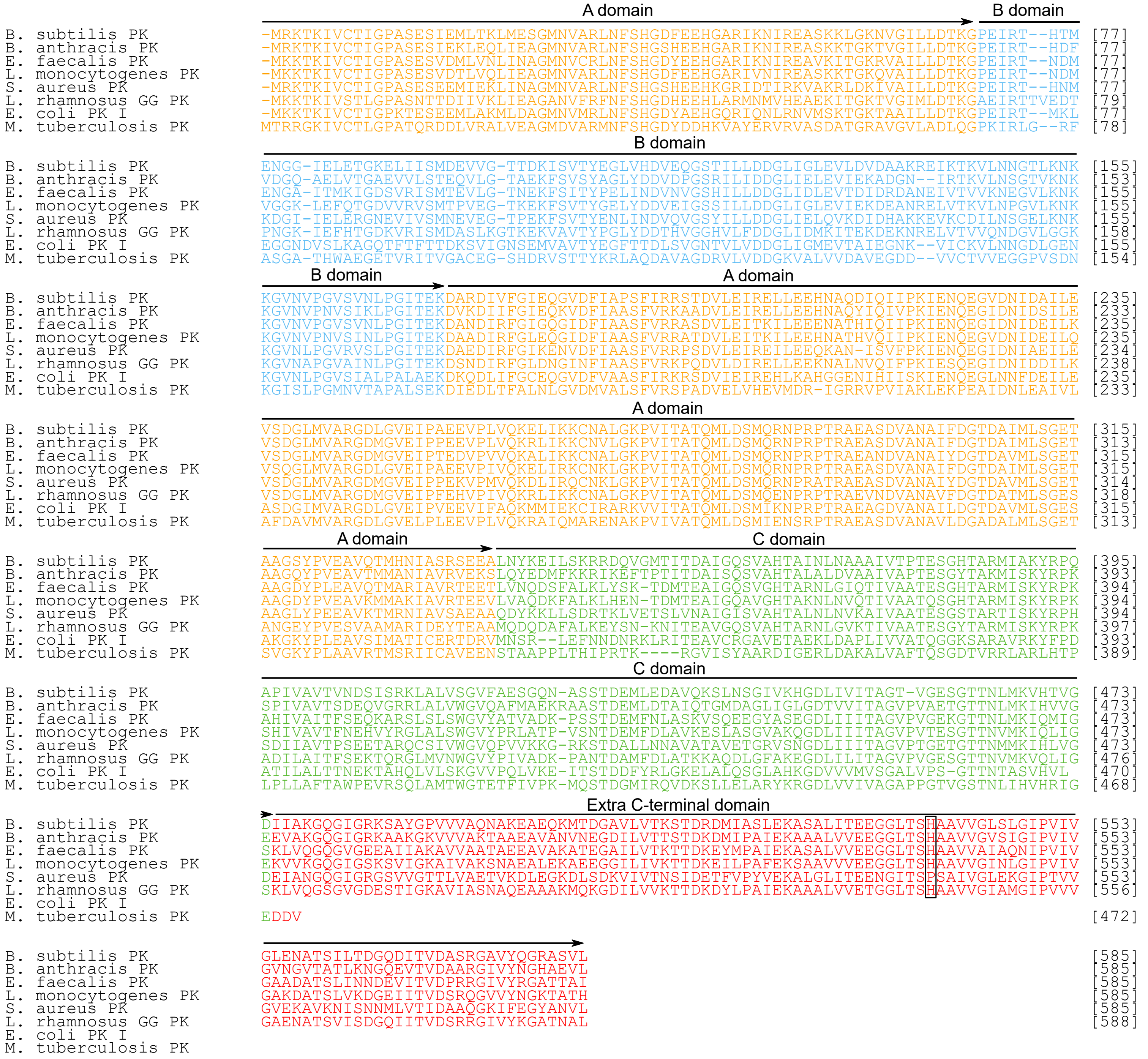


**Supplementary Fig. 1: Pyruvate kinases in some bacteria contain an extra C-terminal domain.**

Amino acid sequences alignment of pyruvate kinase in different species. Active site is in a pocket between A domain and B domain. C domain is a regulatory domain with effector binding sites. The active histidine on the extra C-terminal domain is circled by a rectangle, which is conserved in most Bacillota species but not *S. aureus*.

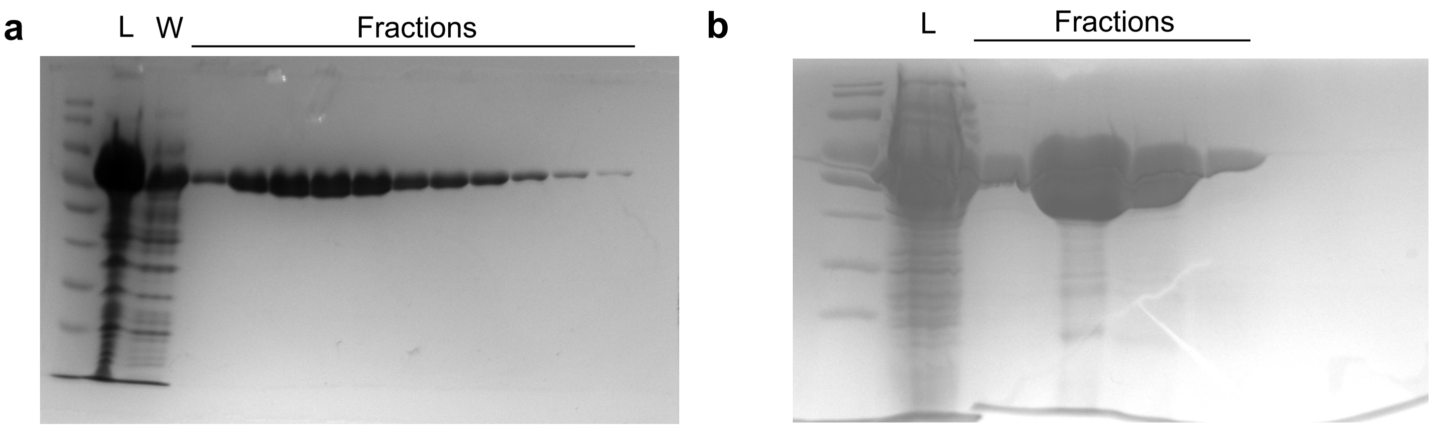


**Supplementary Fig. 2: Purification of *B. subtilis* pyruvate kinases.**

Coomassie brilliant blue stained SDS-PAGE gel of recombinantly expressed (a) wild type *Bacillus subtilis* pyruvate kinase and (b) ΔECTD *Bacillus subtilis* pyruvate kinase, purified with Ni-NTA column. L: lysate; W: wash.

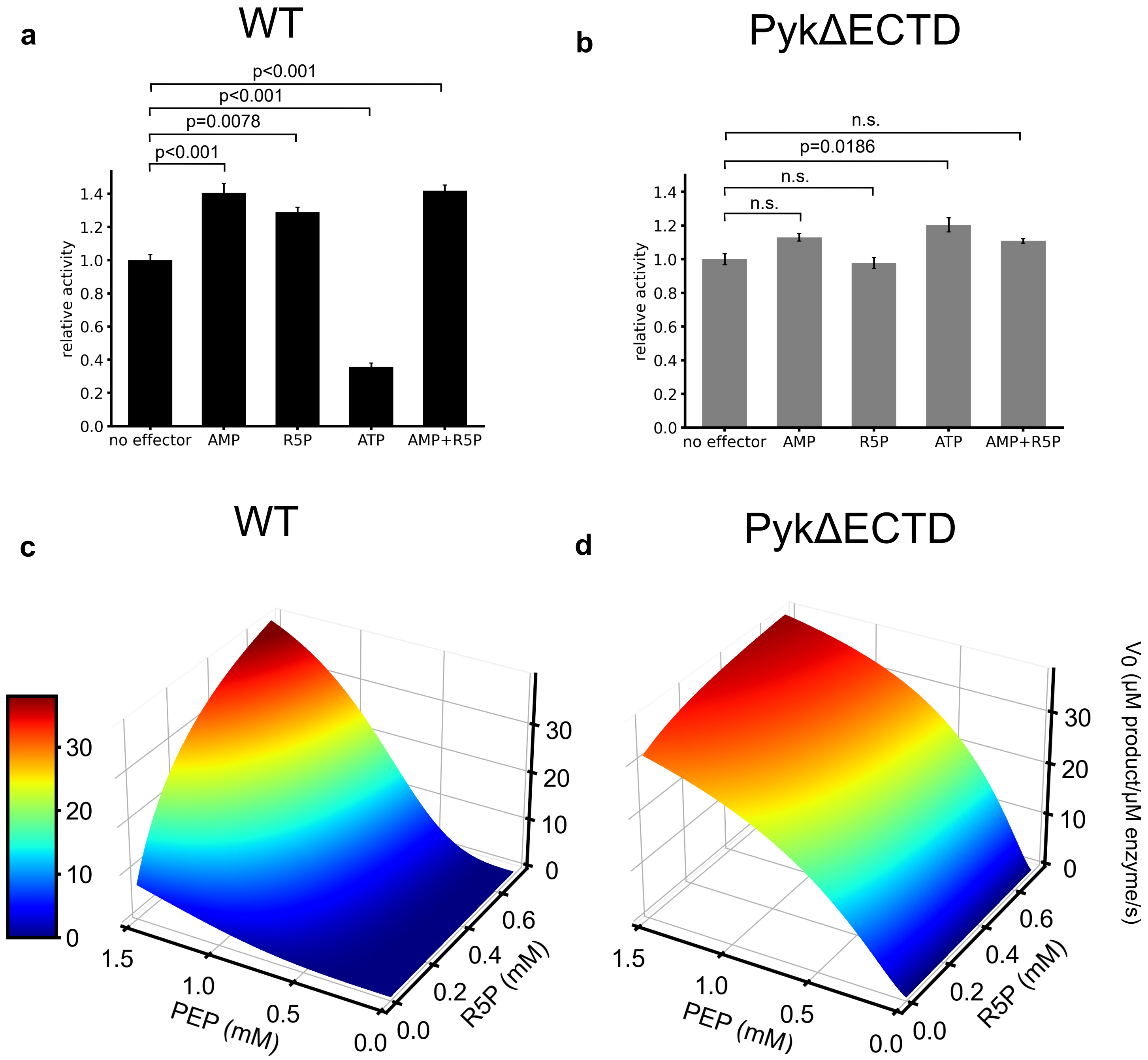


**Supplementary Fig. 3: Three-dimensional analysis of pyruvate kinase kinetics: effects of substrate and activator concentrations on reaction rate.**

(a)-(b) Regulation of allosteric effectors on (a) wild type enzyme and (b) the ΔECTD enzyme variant. (c)-(d) Three-dimensional enzyme kinetic data in different combination of the substate PEP and activator R5P from Fig. 1d and 1e were fitted to an adapted nonessential activation equation[58] by SciPy, and the fitted data were plotted with Matplotlib. Coefficient of determination ($r^2$) was 0.98 for the fitting of wild type data and 0.97 for the fitting of PykΔECTD data. The catalytic rate constant and dissociation constants from the fit are listed in Table 1.

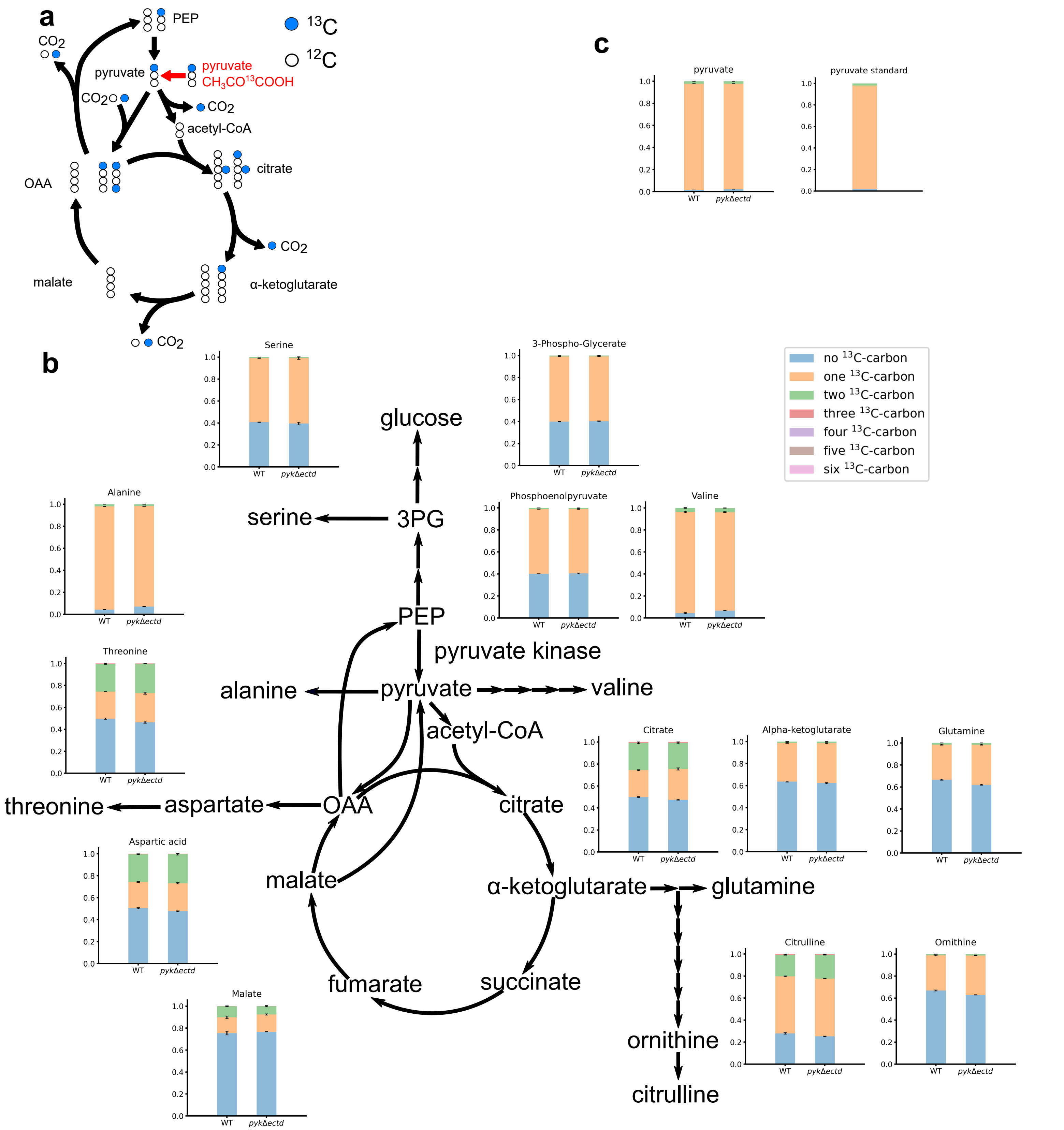


**Supplementary Fig. 4: Isotope composition of important metabolites in cells grown in media with [1-$^{13}$C] pyruvate.**(a) Schematics of the labeling experiment using [1-$^{13}$C] pyruvate in Fig. 2. In *B. subtilis*, PckA, which converts OAA to PEP, is the only enzyme that regenerates PEP from TCA cycle intermediates. Therefore, when pyruvate is the sole carbon source, all PEP is produced from OAA. OAA can be synthesized from pyruvate directly or from malate. When cells are grown on [1-$^{13}$C] pyruvate, OAA directly synthesized from pyruvate is [1-$^{13}$C] labeled. However, OAA synthesized from malate after one round of the TCA cycle is unlabeled. Therefore, only a fraction of OAA is expected to be labeled, as well as PEP since it is derived from OAA. (b) Wild type and *pykΔectd* cells were grown in media with [1-$^{13}$C] pyruvate as sole carbon source, isotope composition of intracellular metabolites was analyzed by LC-MS. Isotope composition of some metabolites in gluconeogenesis and TCA cycle, and some amino acids are shown here. (c) Isotope composition of pyruvate in cells (left) and in the media (right).

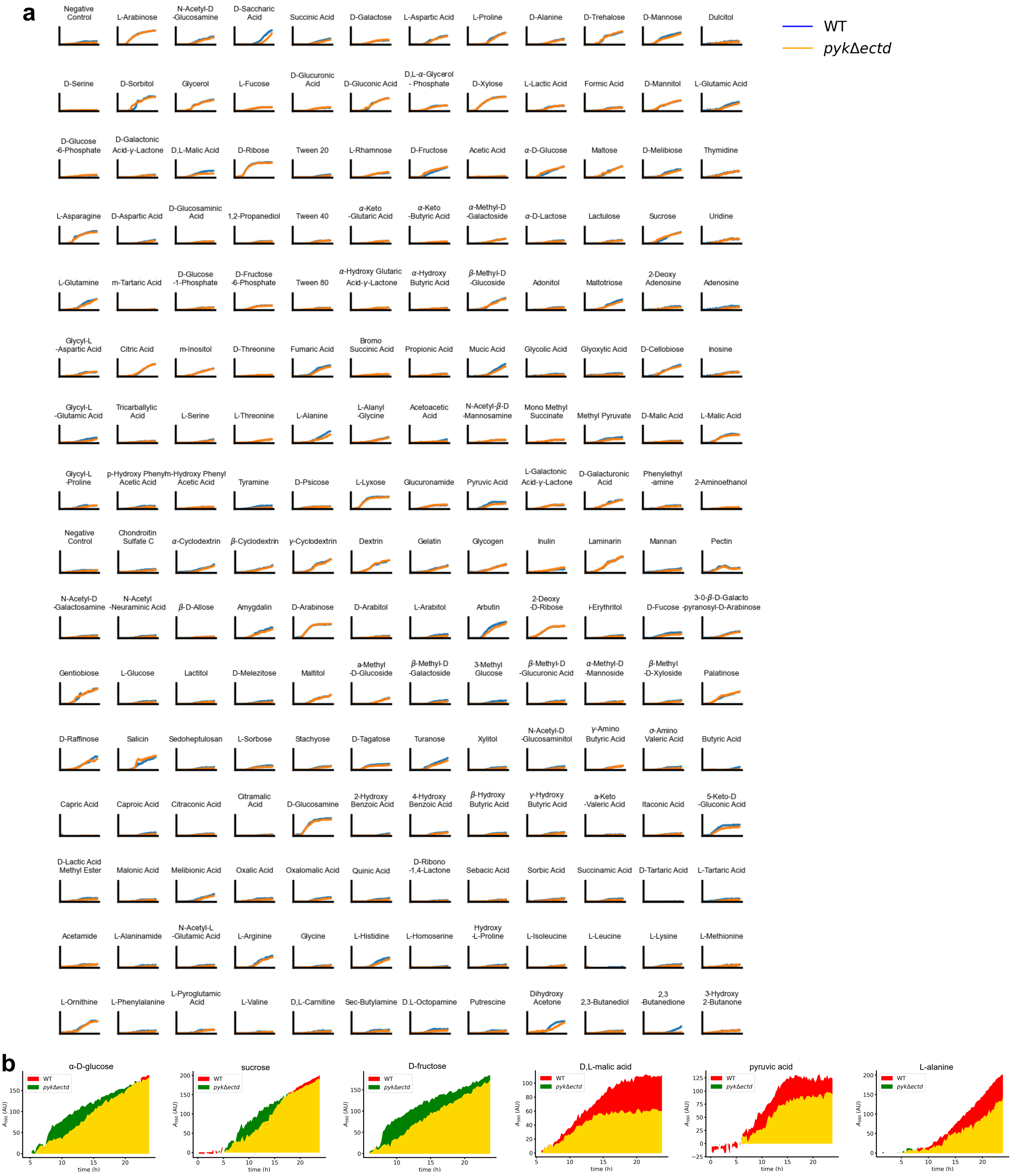


**Supplementary Fig. 5: Phenotype profiling for *pyk*Δectd mutant.**

(a) Cells were inoculated in media with the indicated carbon sources and a redox dye which absorbs at 590 nm when reduced by cellular NADH as a reporter of growth. (b) Examples of several glycolytic (glucose, sucrose and fructose) and gluconeogenic carbon sources (malate, pyruvate and alanine) were plotted separately. Area under wild type curve is colored red, area under *pyk*Δ*ectd* curve is colored green, overlapping region is colored yellow.

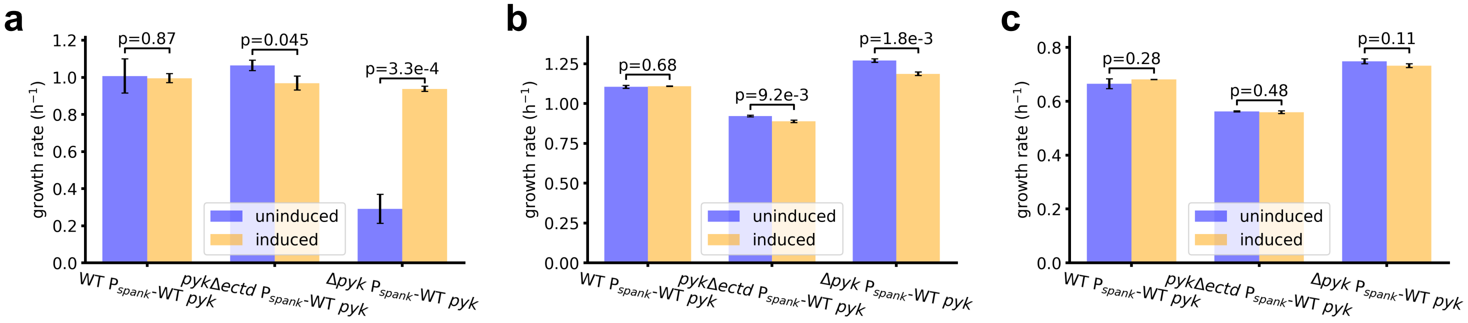


**Supplementary Fig. 6: Complementation tests of *pyk* mutant alleles.**
Growth rates of wild type *pyk*, the *pyk*Δ*ectd* mutant, or the *Δpyk* mutant alleles complemented with wild type *pyk* growing in glycolytic (glucose, a) or gluconeogenic (malate, b or pyruvate, c) media. Wild type *pyk* was provided ectopically at the *amyE* locus with an IPTG-inducible promoter.

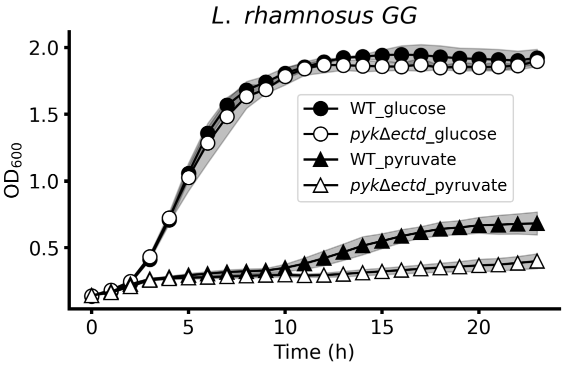


**Supplementary Fig. 7: Growth curves of *L. rhamnosus GG* in media with glycolytic or gluconeogenic carbon sources.**
Growth curve of the *L. rhamnosus GG* wild type and *pyk*Δ*ectd* cells in media with glucose or pyruvate as the sole carbon source. The same data as Fig. 2i.

**a**

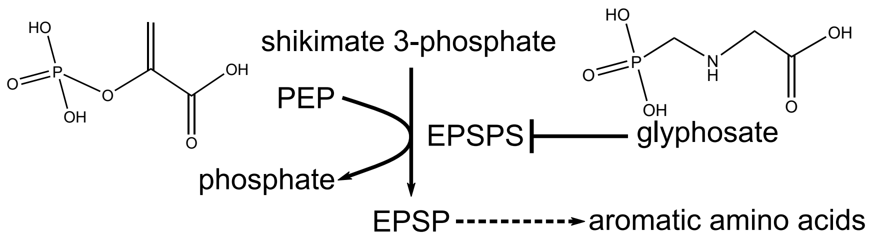


**b**

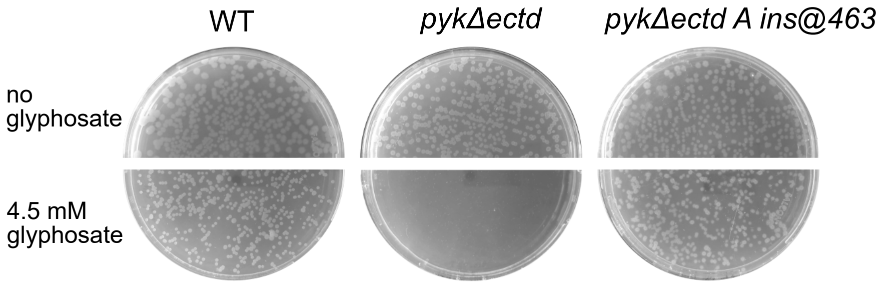


**Supplementary Fig. 8: Hypersensitivity of *pyk*Δectd mutant to glyphosate can be suppressed by a loss-of-function mutation of *pyk*.**

(a) Glyphosate inhibits the shikimate pathway which is required for the synthesis of aromatic amino acids by competing with PEP. EPSP: 5-enolpyruvylshikimate 3-phosphate; EPSPS: EPSP synthase. (b) Plate images of wild type, *pyk*Δ*ectd*, and glyphosate suppressor on tmalate minimal plates with or without glyphosate. The same data as Fig. 2l.

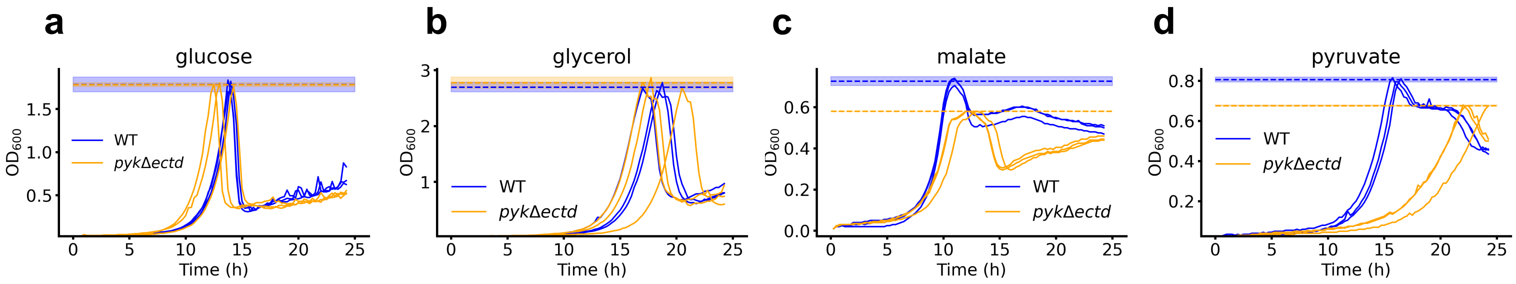


**Supplementary Fig. 9: Maximum $OD_{600}$ of wild type or mutant cells grown in media with different sole carbon sources.** Growth curve of wild type or *pyk*Δ*ectd* cells in media with 2 g/L (a) glucose, (b) glycerol, (c) malate or (d) pyruvate as the sole carbon source, dash lines represent the average maximum ODs for each condition, colored region represents 95% confidence interval.

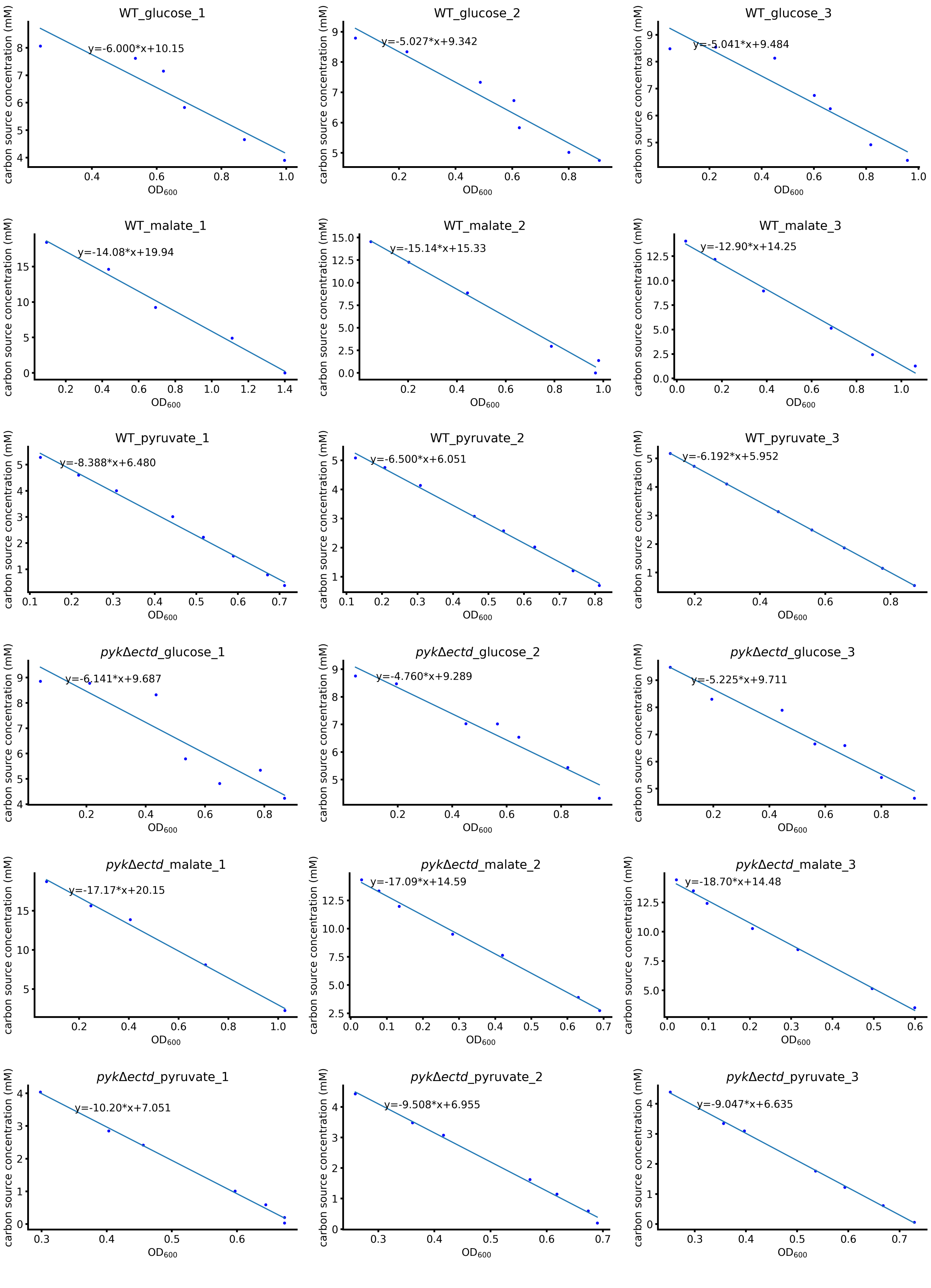


**Supplementary Fig. 10: Gluconeogenic carbon use efficiency of *pyk*Δectd mutant is lower than wild type strain.**

Carbon sources remaining in spent media plotted versus $OD_{600}$ of cells. Slopes are calculated by linear regression with the method of least squares, which represents carbon use efficiency.

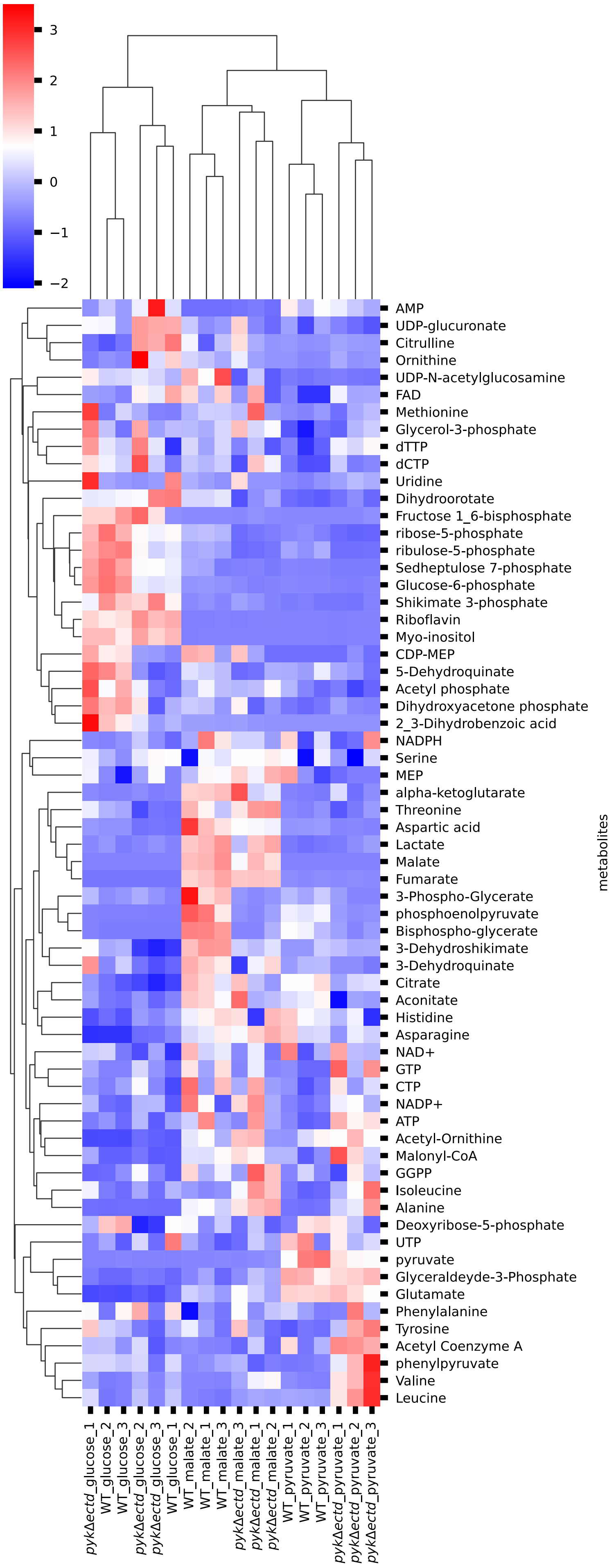


**Supplementary Fig. 11: Metabolome of wild type and *pykΔectd* mutant grown in glycolytic or gluconeogenic media.**
Heatmap of metabolites of wild type and *pykΔectd* cells grown in media with different carbon sources. Rows and columns are clustered with UPGMA (unweighted pair group method with arithmetic mean). Color represents Z scores for rows. MEP: 2-C-methyl-D-erythritol 4-phosphate; GGPP: geranylgeranyl pyrophosphate.

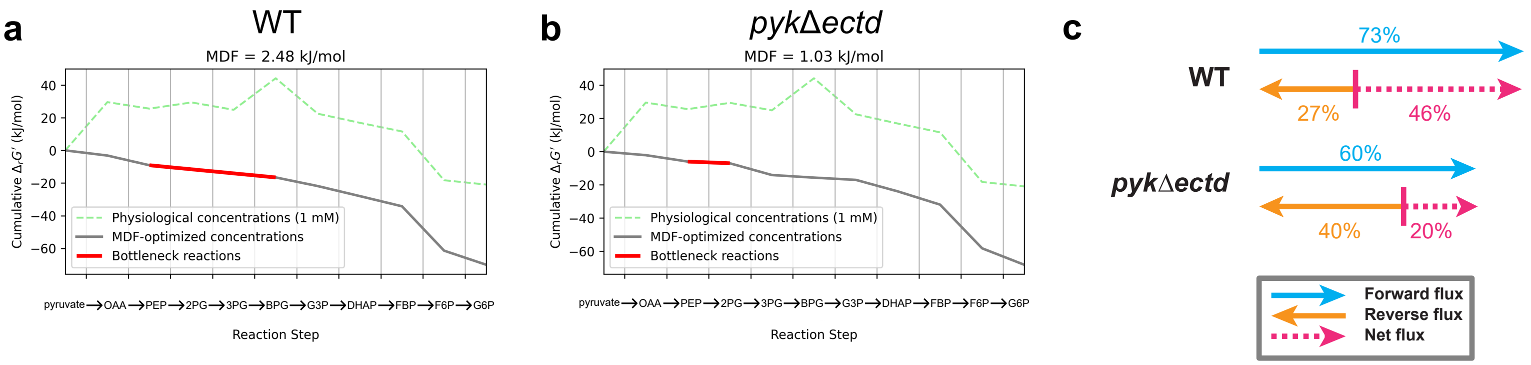


**Supplementary Fig. 12: Thermodynamic analysis of gluconeogenesis for wild type and *pyk*Δ*ectd* cells grown in malate.**
(A)-(B) Cumulative drops in Gibbs free energies of gluconeogenesis for (a) wild type and (b) *pyk*Δ*ectd* cells. MDF: max-min driving force. (c) The calculated forward flux ration, reverse flux ratio and net flux ratio of the bottleneck reaction in gluconeogenesis for wild type and *pyk*Δ*ectd* cells grown in malate.

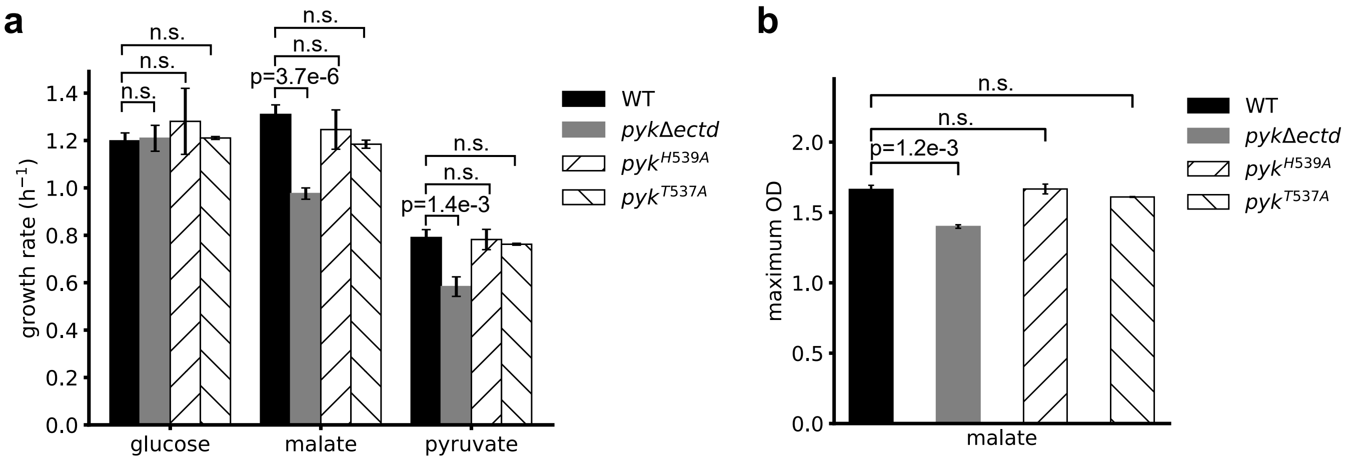


**Supplementary Fig. 13: Potential phosphorylation sites on pyruvate kinase ECTD are not responsible for defective gluconeogenic growth and low growth yield.**

(a) Growth rate of wild type, *pyk*Δ*ectd*, *pyk*$^{H539A}$ and *pyk*$^{T537A}$ cells in media with different sole carbon sources. (b) Maximum $OD_{600}$ of wild type, *pyk*Δ*ectd*, *pyk*$^{H539A}$ and *pyk*$^{T537A}$ cells in media with malate as the sole carbon source.

Supplementary Table 1: Distribution of pyruvate kinase ECTD in the 118 bacterial reference genomes

| Accession number | taxonomy | with pyk ECTD? |
|---|---|---|
| U00096 | Bacteria_Proteobacteria_Gammaproteobacteria_Enterobacterales_Enterobacteriaceae_Escherichia | no |
| AE003852 | Bacteria_Proteobacteria_Gammaproteobacteria_Vibrionales_Vibrionaceae_Vibrio | no |
| AE004091 | Bacteria_Proteobacteria_Gammaproteobacteria_Pseudomonadales_Pseudomonadaceae_Pseudomonas | no |
| AE004092 | Bacteria_Firmicutes_Bacilli_Lactobacillales_Streptococcaceae_Streptococcus | no |
| AE004969 | Bacteria_Proteobacteria_Betaproteobacteria_Neisseriales_Neisseriaceae_Neisseria | no |
| AE005176 | Bacteria_Firmicutes_Bacilli_Lactobacillales_Streptococcaceae_Lactococcus | no |
| AE005673 | Bacteria_Proteobacteria_Alphaproteobacteria_Caulobacterales_Caulobacteraceae_Caulobacter | no |
| AE005674 | Bacteria_Proteobacteria_Gammaproteobacteria_Enterobacterales_Enterobacteriaceae_Shigella | no |
| AE006468 | Bacteria_Proteobacteria_Gammaproteobacteria_Enterobacterales_Enterobacteriaceae_Salmonella | no |
| AL591688 | Bacteria_Proteobacteria_Alphaproteobacteria_Rhizobiales_Rhizobiaceae_Sinorhizobium/Ensifer group_Sinorhizobium | no |
| AE006470 | Bacteria_Chlorobi_Chlorobia_Chlorobiales_Chlorobiaceae_Chlorobaculum | no pyk |
| AE007317 | Bacteria_Firmicutes_Bacilli_Lactobacillales_Streptococcaceae_Streptococcus | no |
| AE008922 | Bacteria_Proteobacteria_Gammaproteobacteria_Xanthomonadales_Xanthomonadaceae_Xanthomonas | no |
| AE009948 | Bacteria_Firmicutes_Bacilli_Lactobacillales_Streptococcaceae_Streptococcus | no |
| AE009951 | Bacteria_Fusobacteria_Fusobacteriales_Fusobacteriaceae_Fusobacterium | no |
| AE014133 | Bacteria_Firmicutes_Bacilli_Lactobacillales_Streptococcaceae_Streptococcus | no |
| AE014295 | Bacteria_Actinobacteria_Bifidobacteriales_Bifidobacteriaceae_Bifidobacterium | no |
| AE015451 | Bacteria_Proteobacteria_Gammaproteobacteria_Pseudomonadales_Pseudomonadaceae_Pseudomonas | no |
| AE015929 | Bacteria_Firmicutes_Bacilli_Bacillales_Staphylococcaceae_Staphylococcus | with pyk ECTD |
| AE016828 | Bacteria_Proteobacteria_Gammaproteobacteria_Legionellales_Coxiellaceae_Coxiella | no |
| AE016830 | Bacteria_Firmicutes_Bacilli_Lactobacillales_Enterococcaceae_Enterococcus | with pyk ECTD |
| AE016853 | Bacteria_Proteobacteria_Gammaproteobacteria_Pseudomonadales_Pseudomonadaceae_Pseudomonas | no |
| AE016877 | Bacteria_Firmicutes_Bacilli_Bacillales_Bacillaceae_Bacillus_Bacillus cereus group | with pyk ECTD |
| AE016879 | Bacteria_Firmicutes_Bacilli_Bacillales_Bacillaceae_Bacillus_Bacillus cereus group | with pyk ECTD |
| AE017126 | Bacteria_Cyanobacteria_Synechococcales_Prochloraceae_Prochlorococcus | with pyk ECTD |
| AE017180 | Bacteria_Proteobacteria_Deltaproteobacteria_Desulfuromonadales_Geobacteraceae_Geobacter | no |
| AE017225 | Bacteria_Firmicutes_Bacilli_Bacillales_Bacillaceae_Bacillus_Bacillus cereus group | with pyk ECTD |
| AE017226 | Bacteria_Spirochaetes_Spirochaetales_Spirochaetaceae_Treponema | no |
| AE017263 | Bacteria_Tenericutes_Mollicutes_Entomoplasmatales_Entomoplasmataceae_Mesoplasma | no |
| AE017354 | Bacteria_Proteobacteria_Gammaproteobacteria_Legionellales_Legionellaceae_Legionella | no |
| AE017355 | Bacteria_Firmicutes_Bacilli_Bacillales_Bacillaceae_Bacillus_Bacillus cereus group | with pyk ECTD |
| AE000511 | Bacteria_Proteobacteria_Epsilonproteobacteria_Campylobacterales_Helicobacteraceae_Helicobacter | no pyk |
| AE000512 | Bacteria_Thermotogae_Thermotogales_Thermotogaceae_Thermotoga | no |
| AE000513 | Bacteria_Deinococcus-Thermus_Deinococci_Deinococcales_Deinococcaceae_Deinococcus | no |
| AE000657 | Bacteria_Aquificae_Aquificales_Aquificaceae_Aquifex | no pyk |
| AE000783 | Bacteria_Spirochaetes_Spirochaetales_Borreliaceae_Borreliella | no |
| AE001273 | Bacteria_Chlamydiae_Chlamydiales_Chlamydiaceae_Chlamydia/Chlamydophila group_Chlamydia | no |
| AE001363 | Bacteria_Chlamydiae_Chlamydiales_Chlamydiaceae_Chlamydia/Chlamydophila group_Chlamydia | no |
| AE001437 | Bacteria_Firmicutes_Clostridia_Clostridiales_Clostridiaceae_Clostridium | no |
| AE002098 | Bacteria_Proteobacteria_Betaproteobacteria_Neisseriales_Neisseriaceae_Neisseria | no |
| BA000007 | Bacteria_Proteobacteria_Gammaproteobacteria_Enterobacterales_Enterobacteriaceae_Escherichia | no |
| CP000233 | Bacteria_Firmicutes_Bacilli_Lactobacillales_Lactobacillaceae_Lactobacillus | with pyk ECTD |
| AJ749949 | Bacteria_Proteobacteria_Gammaproteobacteria_Thiotrichales_Francisellaceae_Francisella | no |
| AL009126 | Bacteria_Firmicutes_Bacilli_Bacillales_Bacillaceae_Bacillus | with pyk ECTD |
| AL590842 | Bacteria_Proteobacteria_Gammaproteobacteria_Enterobacterales_Yersiniaceae_Yersinia | no |
| AL111168 | Bacteria_Proteobacteria_Epsilonproteobacteria_Campylobacterales_Campylobacteraceae_Campylobacter | no |
| AM180355 | Bacteria_Firmicutes_Clostridia_Clostridiales_Peptostreptococcaceae_Clostridioides | with pyk ECTD |
| AM286415 | Bacteria_Proteobacteria_Gammaproteobacteria_Enterobacterales_Yersiniaceae_Yersinia | no |
| BA000003 | Bacteria_Proteobacteria_Gammaproteobacteria_Enterobacterales_Erwiniaceae_Buchnera | no |
| AP006841 | Bacteria_Bacteroidetes_Bacteroidia_Bacteroidales_Bacteroidaceae_Bacteroides | no |
| AE015928 | Bacteria_Bacteroidetes_Bacteroidia_Bacteroidales_Bacteroidaceae_Bacteroides | no |
| BA000036 | Bacteria_Actinobacteria_Corynebacteriales_Corynebacteriaceae_Corynebacterium | no |
| BA000039 | Bacteria_Cyanobacteria_Synechococcales_Synechococcaceae_Thermosynechococcus | with pyk ECTD |
| BA000040 | Bacteria_Proteobacteria_Alphaproteobacteria_Rhizobiales_Bradyrhizobiaceae_Bradyrhizobium | no |
| BA000045 | Bacteria_Cyanobacteria_Gloeobacteria_Gloeobacterales_Gloeobacteraceae_Gloeobacter | with pyk ECTD |
| BX293980 | Bacteria_Tenericutes_Mollicutes_Mycoplasmataceae_Mycoplasma | no |
| BX571965 | Bacteria_Proteobacteria_Betaproteobacteria_Burkholderiales_Burkholderiaceae_Burkholderia_pseudomallei group | no |
| CP000010 | Bacteria_Proteobacteria_Betaproteobacteria_Burkholderiales_Burkholderiaceae_Burkholderia_pseudomallei group | no |
| CP000020 | Bacteria_Proteobacteria_Gammaproteobacteria_Vibrionales_Vibrionaceae_Aliivibrio | no |
| CP000033 | Bacteria_Firmicutes_Bacilli_Lactobacillales_Lactobacillaceae_Lactobacillus | with pyk ECTD |
| CP000034 | Bacteria_Proteobacteria_Gammaproteobacteria_Enterobacterales_Enterobacteriaceae_Shigella | no |
| CP000075 | Bacteria_Proteobacteria_Gammaproteobacteria_Pseudomonadales_Pseudomonadaceae_Pseudomonas_Pseudomonas syringae | no |
| CP000159 | Bacteria_Bacteroidetes_Bacteroidetes Order II. Incertae sedis_Rhodothermaceae_Salinibacter | no |
| CP000230 | Bacteria_Proteobacteria_Alphaproteobacteria_Rhodospirillales_Rhodospirillaceae_Rhodospirillum | no |
| CP000232 | Bacteria_Firmicutes_Clostridia_Thermoanaerobacterales_Thermoanaerobacteraceae_Moorella group_Moorella | with pyk ECTD |
| CP000253 | Bacteria_Firmicutes_Bacilli_Bacillales_Staphylococcaceae_Staphylococcus | with pyk ECTD |
| CP000387 | Bacteria_Firmicutes_Bacilli_Lactobacillales_Streptococcaceae_Streptococcus | no |
| CP000423 | Bacteria_Firmicutes_Bacilli_Lactobacillales_Lactobacillaceae_Lactobacillus | with pyk ECTD |
| CP000462 | Bacteria_Proteobacteria_Gammaproteobacteria_Aeromonadales_Aeromonadaceae_Aeromonas | no |
| CP000480 | Bacteria_Actinobacteria_Corynebacteriales_Mycobacteriaceae_Mycobacterium | no |
| CP000727 | Bacteria_Firmicutes_Clostridia_Clostridiales_Clostridiaceae_Clostridium | with pyk ECTD |
| CP000738 | Bacteria_Proteobacteria_Alphaproteobacteria_Rhizobiales_Rhizobiaceae_Sinorhizobium/Ensifer group_Sinorhizobium | no |
| CP001389 | Bacteria_Proteobacteria_Alphaproteobacteria_Rhizobiales_Rhizobiaceae_Sinorhizobium/Ensifer group_Sinorhizobium | no |
| CP000909 | Bacteria_Chloroflexi_Chloroflexia_Chloroflexales_Chloroflexineae_Chloroflexaceae_Chloroflexus | no |
| CP001147 | Bacteria_Nitrospirae_Nitrospirales_Nitrospiraceae_Thermodesulfovibrio | no |
| CP001251 | Bacteria_Dictyoglomi_Dictyoglomales_Dictyoglomaceae_Dictyoglomus | with pyk ECTD |
| CP001340 | Bacteria_Proteobacteria_Alphaproteobacteria_Caulobacterales_Caulobacteraceae_Caulobacter | no |
| CP001643 | Bacteria_Actinobacteria_Micrococcales_Dermabacteraceae_Brachybacterium | no |
| CP001818 | Bacteria_Synergistetes_Synergistia_Synergistales_Synergistaceae_Thermanaerovibrio | with pyk ECTD |
| CP001918 | Bacteria_Proteobacteria_Gammaproteobacteria_Enterobacterales_Enterobacteriaceae_Enterobacter_Enterobacter cloacae complex | no |
| CU928164 | Bacteria_Proteobacteria_Gammaproteobacteria_Enterobacterales_Enterobacteriaceae_Escherichia | no |
| FM252032 | Bacteria_Firmicutes_Bacilli_Lactobacillales_Streptococcaceae_Streptococcus | no |
| FN568063 | Bacteria_Firmicutes_Bacilli_Lactobacillales_Streptococcaceae_Streptococcus | no |
| L42023 | Bacteria_Proteobacteria_Gammaproteobacteria_Pasteurellales_Pasteurellaceae_Haemophilus | no |
| U00089 | Bacteria_Tenericutes_Mollicutes_Mycoplasmataceae_Mycoplasma | no |
| AM412317 | Bacteria_Firmicutes_Clostridia_Clostridiales_Clostridiaceae_Clostridium | with pyk ECTD |
| AM398681 | Bacteria_Bacteroidetes_Flavobacteriia_Flavobacteriales_Flavobacteriaceae_Flavobacterium | no |
| AM884176 | Bacteria_Chlamydiae_Chlamydiales_Chlamydiaceae_Chlamydia/Chlamydophila group_Chlamydia | no |
| CU458896 | Bacteria_Actinobacteria_Corynebacteriales_Mycobacteriaceae_Mycobacteroides abscessus | no |
| AP008226 | Bacteria_Deinococcus-Thermus_Deinococci_Thermales_Thermaceae_Thermus | no |
| AE007869 | Bacteria_Proteobacteria_Alphaproteobacteria_Rhizobiales_Rhizobiaceae_Rhizobium/Agrobacterium group_Agrobacterium_Agrobacterium tumefaciens complex | no |

| | | |
|---|---|---|
| AE010300 | Bacteria_Spirochaetes_Leptospirales_Leptospiraceae_Leptospira | no |
| AE014299 | Bacteria_Proteobacteria_Gammaproteobacteria_Alteromonadales_Shewanellaceae_Shewanella | no |
| CP002104 | Bacteria_Actinobacteria_Bifidobacteriales_Bifidobacteriaceae_Gardnerella | no |
| CP001840 | Bacteria_Actinobacteria_Bifidobacteriales_Bifidobacteriaceae_Bifidobacterium | no |
| CP003583 | Bacteria_Firmicutes_Bacilli_Lactobacillales_Enterococcaceae_Enterococcus | with pyk ECTD |
| CP001855 | Bacteria_Proteobacteria_Gammaproteobacteria_Enterobacterales_Enterobacteriaceae_Escherichia | no |
| CP002447 | Bacteria_Proteobacteria_Alphaproteobacteria_Rhizobiales_Phyllobacteriaceae_Mesorhizobium | no |
| CP002177 | Bacteria_Proteobacteria_Gammaproteobacteria_Pseudomonadales_Moraxellaceae_Acinetobacter_Acinetobacter calcoaceticus/baumannii complex | no pyk |
| BX470248 | Bacteria_Proteobacteria_Betaproteobacteria_Burkholderiales_Alcaligenaceae_Bordetella | no |
| AJ235269 | Bacteria_Proteobacteria_Alphaproteobacteria_Rickettsiales_Rickettsiaceae_Rickettsieae_Rickettsia_typhus group | no pyk |
| AE017285 | Bacteria_Proteobacteria_Deltaproteobacteria_Desulfovibrionales_Desulfovibrionaceae_Desulfovibrio | no |
| LT708304 | Bacteria_Actinobacteria_Corynebacteriales_Mycobacteriaceae_Mycobacterium_Mycobacterium tuberculosis complex | no |
| AL450380 | Bacteria_Actinobacteria_Corynebacteriales_Mycobacteriaceae_Mycobacterium | no |
| AL123456 | Bacteria_Actinobacteria_Corynebacteriales_Mycobacteriaceae_Mycobacterium_Mycobacterium tuberculosis complex | no |
| AL513382 | Bacteria_Proteobacteria_Gammaproteobacteria_Enterobacterales_Enterobacteriaceae_Salmonella | no |
| AL591824 | Bacteria_Firmicutes_Bacilli_Bacillales_Listeriaceae_Listeria | with pyk ECTD |
| BA000031 | Bacteria_Proteobacteria_Gammaproteobacteria_Vibrionales_Vibrionaceae_Vibrio | no |
| BX119912 | Bacteria_Planctomycetes_Planctomycetia_Planctomycetales_Planctomycetaceae_Rhodopirellula | no |
| CP002000 | Bacteria_Actinobacteria_Pseudonocardiales_Pseudonocardiaceae_Amycolatopsis | no |
| AL645882 | Bacteria_Actinobacteria_Streptomycetales_Streptomycetaceae_Streptomyces_Streptomyces albidoflavus group | no |
| AL935263 | Bacteria_Firmicutes_Bacilli_Lactobacillales_Lactobacillaceae_Lactobacillus | with pyk ECTD |
| CP002824 | Bacteria_Proteobacteria_Gammaproteobacteria_Enterobacterales_Enterobacteriaceae_Klebsiella | no |
| CP002018 | Bacteria_Proteobacteria_Alphaproteobacteria_Rhodobacterales_Rhodobacteraceae_Ketogulonicigenium | no |
| CP003200 | Bacteria_Proteobacteria_Gammaproteobacteria_Enterobacterales_Enterobacteriaceae_Klebsiella | no |
| CP003289 | Bacteria_Proteobacteria_Gammaproteobacteria_Enterobacterales_Enterobacteriaceae_Escherichia | no |
| HE965803 | Bacteria_Proteobacteria_Betaproteobacteria_Burkholderiales_Alcaligenaceae_Bordetella | no |
| HE965806 | Bacteria_Proteobacteria_Betaproteobacteria_Burkholderiales_Alcaligenaceae_Bordetella | no |

Supplementary Table 2: Relative abundance of metabolites in wild type and *pykΔectd* cells, measured by spiking in 13C labeled metabolites from wild type cells grown in [U-13C] glucose

| strain | | WT | | | | | | | | | pykΔectd | | | | | | | | |
|---|---|---|---|---|---|---|---|---|---|---|---|---|---|---|---|---|---|---|---|
| carbon source | | glucose | | | malate | | | pyruvate | | | glucose | | | malate | | | pyruvate | | |
| repeat | | 1 | 2 | 3 | 1 | 2 | 3 | 1 | 2 | 3 | 1 | 2 | 3 | 1 | 2 | 3 | 1 | 2 | 3 |
| | normalization | | | | | | | | | | | | | | | | | | |
| Valine | normalized to 13C labeled internal control | 4.24E+00 | 1.83E+00 | 1.97E+00 | 2.21E+00 | 2.23E+00 | 2.11E+00 | 2.48E+00 | 2.07E+00 | 2.00E+00 | 3.37E+00 | 4.05E+00 | 2.23E+00 | 6.60E+00 | 7.44E+00 | 3.49E+00 | 8.20E+00 | 1.00E+01 | 1.56E+01 |
| UTP | normalized to 13C labeled internal control | 2.27E+01 | 1.19E+01 | 8.37E+00 | 1.01E+01 | 1.22E+01 | 8.12E+00 | 1.92E+01 | 2.21E+01 | 9.80E+00 | 1.01E+01 | 1.42E+01 | 9.03E+00 | 1.36E+01 | 1.15E+01 | 9.00E+00 | 1.70E+01 | 1.27E+01 | 1.44E+01 |
| Uridine | normalized to 13C labeled internal control | 2.83E+01 | 6.71E+00 | 5.61E+00 | 6.21E+00 | 7.87E+00 | 4.73E+00 | 4.21E+00 | 3.07E+00 | 4.92E+00 | 3.75E+01 | 4.77E+00 | 5.78E+00 | 5.09E+00 | 5.07E+00 | 1.99E+01 | 6.36E+00 | 9.05E+00 | 7.58E+00 |
| UDP-N-acetylglucosamine | normalized to 13C labeled internal control | 2.11E+00 | 2.86E+00 | 3.02E+00 | 4.09E+00 | 5.92E+00 | 8.45E+00 | 7.35E-01 | 5.41E-01 | 7.03E-01 | 4.46E+00 | 3.45E+00 | 2.95E+00 | 2.65E+00 | 0.00E+00 | 0.00E+00 | 7.95E-01 | 6.69E-01 | 5.66E-01 |
| UDP-glucuronate | normalized to 13C labeled internal control | 3.23E+00 | 2.39E+00 | 1.59E+00 | 1.45E+00 | 1.98E+00 | 1.58E+00 | 1.63E+00 | 8.58E-01 | 1.67E+00 | 2.42E+00 | 3.35E+00 | 3.25E+00 | 1.42E+00 | 1.15E+00 | 2.90E+00 | 1.36E+00 | 1.18E+00 | 9.75E-01 |
| Tyrosine | normalized to 13C labeled internal control | 2.15E+00 | 2.38E+00 | 2.25E+00 | 2.03E+00 | 2.56E+00 | 1.72E+00 | 1.52E+00 | 1.73E+00 | 1.67E+00 | 3.09E+00 | 1.77E+00 | 1.58E+00 | 2.01E+00 | 1.66E+00 | 3.13E+00 | 2.27E+00 | 3.31E+00 | 3.63E+00 |
| Threonine | normalized to 13C labeled internal control | 1.07E+00 | 1.35E+00 | 1.29E+00 | 1.69E+00 | 1.96E+00 | 1.42E+00 | 1.18E+00 | 1.15E+00 | 1.22E+00 | 1.57E+00 | 9.28E-01 | 1.09E+00 | 2.06E+00 | 2.08E+00 | 1.76E+00 | 9.78E-01 | 1.11E+00 | 1.25E+00 |
| Shikimate 3-phosphate | normalized to 13C labeled internal control | 3.74E+00 | 6.17E+00 | 4.77E+00 | 7.22E-01 | 5.95E-01 | 4.94E-01 | 2.42E-01 | 3.35E-01 | 7.05E-02 | 3.12E+00 | 4.43E+00 | 6.47E+00 | 8.14E-01 | 4.24E-01 | 1.11E+00 | 8.87E-02 | 3.10E-02 | 3.70E-01 |
| ribose-5-phosphate | normalized to 13C labeled internal control | 2.41E+00 | 4.20E+00 | 3.37E+00 | 1.54E+00 | 1.44E+00 | 1.38E+00 | 7.96E-01 | 8.79E-01 | 7.12E-01 | 3.26E+00 | 2.41E+00 | 2.18E+00 | 6.02E-01 | 6.77E-01 | 4.33E-01 | 4.16E-01 | 3.49E-01 | 3.77E-01 |
| ribulose-5-phosphate | normalized to 13C labeled internal control | 2.32E+00 | 4.57E+00 | 4.74E+00 | 1.42E+00 | 1.28E+00 | 1.18E+00 | 1.26E+00 | 9.39E-01 | 1.41E+00 | 3.95E+00 | 2.79E+00 | 1.99E+00 | 4.00E-01 | 5.35E-01 | 3.45E-01 | 4.00E-01 | 4.53E-01 | 4.84E-01 |
| pyruvate | normalized to 13C labeled internal control | 1.24E+01 | 6.07E+00 | 4.51E+01 | 6.20E+01 | 6.25E+01 | 7.35E+01 | 1.25E+03 | 2.60E+03 | 2.71E+03 | 7.47E+00 | 6.94E+00 | 3.26E+01 | 9.36E+01 | 1.50E+02 | 5.81E+01 | 1.59E+03 | 1.35E+03 | 1.25E+03 |
| phosphoenolpyruvate | normalized to 13C labeled internal control | 7.98E-01 | 2.23E+00 | 2.08E+00 | 1.27E+02 | 1.40E+02 | 7.25E+01 | 5.39E+01 | 4.88E+01 | 5.78E+01 | 3.56E+00 | 1.84E+00 | 7.91E-01 | 6.10E+00 | 1.71E+01 | 9.76E+00 | 1.53E+01 | 8.13E+00 | 9.46E+00 |
| Leucine | normalized to 13C labeled internal control | 4.19E+00 | 2.41E+00 | 2.60E+00 | 2.56E+00 | 2.74E+00 | 2.42E+00 | 3.45E+00 | 3.36E+00 | 3.26E+00 | 4.66E+00 | 4.11E+00 | 2.78E+00 | 3.43E+00 | 3.30E+00 | 3.12E+00 | 6.06E+00 | 7.90E+00 | 1.03E+01 |
| Lactate | normalized to 13C labeled internal control | 1.28E-01 | 1.74E-01 | 1.08E-01 | 6.14E-01 | 5.70E-01 | 6.98E-01 | 1.05E-01 | 8.01E-02 | 9.80E-02 | 1.41E-01 | 1.33E-01 | 1.65E-01 | 5.77E-01 | 6.55E-01 | 2.71E-01 | 1.11E-01 | 1.49E-01 | 1.96E-01 |
| Isoleucine | normalized to 13C labeled internal control | 2.86E+00 | 2.26E+00 | 2.07E+00 | 2.38E+00 | 2.57E+00 | 2.46E+00 | 2.24E+00 | 1.96E+00 | 2.03E+00 | 3.49E+00 | 2.68E+00 | 2.13E+00 | 4.75E+00 | 4.25E+00 | 3.55E+00 | 2.81E+00 | 3.67E+00 | 5.14E+00 |
| Glyceraldeyde-3-Phosphate | normalized to OD600 | 1.61E+04 | 4.83E+04 | 3.95E+04 | 3.53E+07 | 1.83E+07 | 2.87E+05 | 1.40E+08 | 1.35E+08 | 9.66E+07 | 1.21E+07 | 1.12E+07 | 8.46E+06 | 5.25E+07 | 3.72E+07 | 2.19E+07 | 1.12E+08 | 1.17E+08 | 1.30E+08 |
| Glutamate | normalized to 13C labeled internal control | 1.79E+00 | 1.50E+00 | 1.49E+00 | 2.48E+00 | 2.65E+00 | 2.19E+00 | 3.51E+00 | 3.46E+00 | 3.51E+00 | 1.45E+00 | 1.64E+00 | 1.50E+00 | 2.64E+00 | 2.28E+00 | 3.10E+00 | 3.69E+00 | 3.41E+00 | 3.11E+00 |
| Dihydroorotate | normalized to 13C labeled internal control | 2.94E+00 | 1.71E+00 | 1.79E+00 | 1.53E+00 | 1.53E+00 | 1.67E+00 | 6.80E-01 | 6.04E-01 | 5.71E-01 | 1.67E+00 | 1.83E+00 | 2.87E+00 | 9.45E-01 | 1.16E+00 | 4.78E-01 | 7.21E-01 | 9.31E-01 | 6.44E-01 |
| Citrulline | normalized to 13C labeled internal control | 2.40E+00 | 7.46E-01 | 9.17E-01 | 7.90E-01 | 1.61E+00 | 1.33E+00 | 1.11E+00 | 1.05E+00 | 1.07E+00 | 9.08E-01 | 2.21E+00 | 2.12E+00 | 1.22E+00 | 1.09E+00 | 1.85E+00 | 1.16E+00 | 1.13E+00 | 1.11E+00 |
| Citrate | normalized to 13C labeled internal control | 2.89E+00 | 3.99E+00 | 3.37E+00 | 7.87E+00 | 8.50E+00 | 6.31E+00 | 7.01E+00 | 6.94E+00 | 7.69E+00 | 4.74E+00 | 2.99E+00 | 2.20E+00 | 5.65E+00 | 4.75E+00 | 8.30E+00 | 4.96E+00 | 6.06E+00 | 6.27E+00 |
| Bisphospho-glycerate | normalized to OD600 | 2.70E+04 | 0.00E+00 | 2.18E+04 | 1.28E+07 | 1.29E+07 | 1.16E+07 | 6.81E+06 | 5.99E+06 | 3.26E+06 | 3.98E+05 | 5.40E+04 | 0.00E+00 | 4.98E+05 | 2.65E+06 | 5.48E+05 | 1.87E+06 | 5.44E+05 | 1.53E+06 |
| ATP | normalized to 13C labeled internal control | 9.93E+00 | 9.17E+00 | 7.16E+00 | 2.02E+01 | 1.21E+01 | 9.96E+00 | 8.63E+00 | 6.63E+00 | 7.05E+00 | 8.00E+00 | 1.09E+01 | 6.56E+00 | 1.95E+01 | 1.03E+01 | 8.59E+00 | 1.82E+01 | 1.50E+01 | 1.60E+01 |
| Aspartic acid | normalized to 13C labeled internal control | 5.71E-01 | 1.55E+00 | 1.58E+00 | 7.20E+00 | 1.12E+01 | 5.93E+00 | 1.68E+00 | 1.75E+00 | 1.86E+00 | 1.81E+00 | 5.98E-01 | 7.21E-01 | 4.83E+00 | 4.56E+00 | 5.04E+00 | 1.21E+00 | 1.27E+00 | 1.08E+00 |
| alpha-ketoglutarate | normalized to 13C labeled internal control | 7.56E-02 | 7.37E-02 | 7.54E-02 | 5.25E-01 | 5.12E-01 | 5.70E-01 | 7.42E-02 | 6.26E-02 | 5.11E-02 | 9.33E-02 | 9.73E-02 | 6.35E-02 | 2.73E-01 | 3.34E-01 | 8.43E-01 | 3.02E-01 | 2.33E-02 | 9.72E-02 |
| Alanine | normalized to OD600 | 1.38E+06 | 1.03E+06 | 2.80E+06 | 9.85E+07 | 9.09E+07 | 6.76E+07 | 6.89E+06 | 4.01E+06 | 6.72E+06 | 3.83E+06 | 2.09E+06 | 7.41E+05 | 1.44E+08 | 1.55E+08 | 1.15E+08 | 6.58E+07 | 8.49E+07 | 1.69E+08 |
| Aconitate | normalized to 13C labeled internal control | 7.50E-01 | 6.88E-01 | 6.42E-01 | 1.66E+00 | 1.74E+00 | 1.40E+00 | 1.23E+00 | 1.32E+00 | 1.48E+00 | 9.02E-01 | 8.50E-01 | 5.87E-01 | 9.89E-01 | 1.07E+00 | 2.23E+00 | 8.45E-02 | 9.21E-01 | 8.80E-01 |
| Acetyl-Ornithine | normalized to 13C labeled internal control | 1.36E+00 | 6.75E-01 | 6.45E-01 | 9.56E+00 | 8.26E+00 | 5.57E+00 | 4.25E+00 | 7.65E+00 | 9.91E+00 | 6.52E-01 | 2.32E+00 | 1.75E+00 | 1.29E+01 | 4.36E+00 | 1.24E+01 | 9.49E+00 | 1.30E+01 | 9.43E+00 |
| Acetyl Coenzyme A | normalized to 13C labeled internal control | 3.41E+00 | 5.55E+00 | 3.79E+00 | 3.35E+00 | 3.75E+00 | 3.22E+00 | 9.12E+00 | 2.85E+00 | 4.99E+00 | 5.54E+00 | 6.44E+00 | 1.75E+00 | 5.91E+00 | 2.35E+00 | 2.39E+00 | 1.21E+01 | 1.16E+01 | 1.08E+01 |
| 5-Dehydroquinate | normalized to 13C labeled internal control | 1.56E+00 | 2.91E+00 | 2.61E+00 | 2.07E+00 | 2.15E+00 | 1.99E+00 | 1.90E+00 | 1.83E+00 | 2.21E+00 | 3.07E+00 | 1.76E+00 | 1.48E+00 | 1.61E+00 | 1.91E+00 | 1.56E+00 | 1.89E+00 | 1.79E+00 | 1.61E+00 |
| 3-Phospho-Glycerate | normalized to OD600 | 2.83E+06 | 1.34E+07 | 2.32E+07 | 1.68E+08 | 3.68E+08 | 1.95E+08 | 5.88E+07 | 3.70E+07 | 7.48E+07 | 5.88E+07 | 4.44E+07 | 2.18E+07 | 1.15E+07 | 2.12E+07 | 2.39E+07 | 3.07E+07 | 7.91E+06 | 7.15E+06 |
| Serine | normalized to 13C labeled internal control | 7.18E+01 | 5.20E+01 | 4.09E+01 | 6.93E+01 | 7.06E+00 | 6.38E+01 | 7.01E+01 | 5.85E+00 | 6.67E+01 | 6.67E+01 | 6.55E+01 | 7.25E+01 | 7.12E+01 | 7.60E+01 | 7.25E+01 | 4.43E+01 | 2.78E+00 | 6.02E+01 |
| Sedheptulose 7-phosphate | normalized to OD600 | 3.08E+07 | 7.32E+07 | 6.06E+07 | 1.09E+07 | 1.29E+07 | 5.10E+05 | 3.02E+06 | 2.05E+06 | 2.38E+06 | 6.17E+07 | 3.52E+07 | 3.66E+07 | 2.43E+06 | 2.97E+04 | 1.48E+06 | 6.87E+04 | 5.06E+05 | 8.41E+04 |
| Riboflavin | normalized to OD600 | 1.39E+07 | 9.80E+06 | 1.06E+07 | 9.56E+04 | 1.14E+05 | 1.60E+05 | 1.80E+05 | 1.49E+05 | 1.24E+05 | 1.14E+07 | 1.57E+07 | 1.31E+07 | 2.54E+05 | 3.51E+05 | 0.00E+00 | 1.20E+05 | 2.41E+05 | 2.69E+05 |
| phenylpyruvate | normalized to 13C labeled internal control | 3.37E+00 | 3.30E+00 | 3.03E+00 | 2.14E+00 | 2.01E+00 | 2.52E+00 | 1.70E+00 | 1.74E+00 | 1.94E+00 | 3.36E+00 | 3.32E+00 | 1.62E+00 | 1.34E+00 | 1.80E+00 | 2.75E+00 | 3.64E+00 | 5.26E+00 | 7.80E+00 |
| Phenylalanine | normalized to 13C labeled internal control | 3.61E+00 | 1.43E+00 | 3.33E+00 | 1.80E+00 | 1.35E-02 | 1.44E+00 | 2.65E+00 | 1.97E+00 | 1.52E+00 | 3.17E+00 | 4.28E+00 | 1.45E+00 | 1.70E+00 | 2.42E+00 | 3.18E+00 | 1.61E+00 | 4.91E+00 | 2.25E+00 |
| Ornithine | normalized to 13C labeled internal control | 2.26E+00 | 1.03E+00 | 9.39E-01 | 1.40E+00 | 1.54E+00 | 1.19E+00 | 1.04E+00 | 9.43E-01 | 9.65E-01 | 8.56E-01 | 3.90E+00 | 1.61E+00 | 1.13E+00 | 1.03E+00 | 1.74E+00 | 1.21E+00 | 1.01E+00 | 1.06E+00 |
| NADPH | normalized to OD600 | 4.21E+05 | 6.99E+05 | 8.45E+05 | 3.32E+06 | 1.09E+06 | 2.23E+06 | 2.44E+06 | 1.09E+05 | 1.63E+06 | 7.45E+05 | 1.41E+06 | 8.21E+05 | 1.51E+06 | 9.88E+05 | 1.52E+06 | 3.36E+05 | 5.04E+05 | 3.15E+06 |
| NADP+ | normalized to OD600 | 1.60E+05 | 1.49E+06 | 8.55E+05 | 1.20E+07 | 2.08E+07 | 0.00E+00 | 2.96E+06 | 1.36E+06 | 2.92E+06 | 6.85E+06 | 6.66E+06 | 6.75E+06 | 1.96E+07 | 5.93E+06 | 1.44E+07 | 1.05E+07 | 1.18E+07 | 6.21E+06 |
| NAD+ | normalized to 13C labeled internal control | 1.26E+00 | 1.83E+00 | 1.52E+00 | 1.82E+00 | 2.23E+00 | 1.90E+00 | 2.43E+00 | 1.37E+00 | 1.70E+00 | 1.80E+00 | 1.36E+00 | 1.66E+00 | 1.91E+00 | 1.50E+00 | 1.57E+00 | 2.30E+00 | 1.75E+00 | 1.72E+00 |
| Myo-inositol | normalized to 13C labeled internal control | 1.37E+00 | 1.36E+00 | 1.00E+00 | 4.23E-03 | 2.60E-03 | 4.36E-03 | 1.05E-02 | 4.10E-03 | 4.21E-03 | 1.35E+00 | 1.52E+00 | 1.18E+00 | 1.27E-02 | 4.91E-03 | 4.28E-03 | 3.13E-03 | 4.12E-03 | 3.87E-03 |
| Methionine | normalized to 13C labeled internal control | 1.14E+00 | 1.05E+00 | 2.64E+00 | 2.51E+00 | 2.03E+00 | 2.48E+00 | 1.38E+00 | 1.24E+00 | 1.59E+00 | 6.77E+00 | 1.97E+00 | 1.20E+00 | 6.10E+00 | 1.67E+00 | 2.20E+00 | 8.69E-01 | 1.88E+00 | 2.22E+00 |
| MEP | normalized to OD600 | 2.98E+07 | 3.02E+07 | 1.96E+07 | 4.15E+07 | 3.50E+07 | 4.04E+07 | 4.95E+07 | 3.10E+07 | 2.40E+07 | 3.94E+07 | 3.24E+07 | 4.05E+07 | 3.97E+07 | 4.79E+07 | 4.49E+07 | 2.76E+07 | 2.92E+07 | 2.93E+07 |
| Malonyl-CoA | normalized to OD600 | 3.45E+05 | 8.13E+05 | 8.36E+05 | 1.28E+07 | 1.29E+07 | 1.64E+07 | 8.80E+06 | 5.01E+05 | 4.89E+06 | 7.57E+06 | 2.71E+06 | 1.14E+05 | 1.68E+07 | 8.72E+06 | 1.92E+07 | 3.24E+07 | 1.99E+07 | 1.10E+07 |
| Malate | normalized to OD600 | 6.63E+05 | 5.15E+05 | 3.52E+05 | 6.63E+11 | 6.41E+11 | 7.90E+11 | 1.82E+06 | 6.92E+05 | 1.22E+06 | 0.00E+00 | 8.55E+05 | 1.40E+07 | 6.50E+11 | 5.28E+11 | 3.93E+11 | 1.12E+07 | 7.56E+06 | 1.15E+07 |
| Histidine | normalized to OD600 | 2.92E+08 | 2.34E+08 | 2.09E+08 | 4.02E+08 | 3.73E+08 | 4.34E+08 | 4.47E+08 | 3.85E+08 | 3.67E+08 | 2.05E+08 | 2.83E+08 | 2.61E+08 | 1.78E+08 | 4.66E+08 | 4.23E+08 | 3.15E+08 | 3.75E+08 | 1.68E+08 |
| GTP | normalized to OD600 | 1.36E+05 | 2.78E+06 | 2.80E+06 | 6.06E+06 | 1.86E+07 | 1.92E+07 | 3.89E+06 | 7.29E+04 | 0.00E+00 | 6.80E+06 | 1.17E+07 | 4.55E+06 | 1.37E+07 | 3.16E+06 | 3.22E+06 | 3.20E+07 | 8.18E+06 | 2.72E+07 |
| Glycerol-3-phosphate | normalized to OD600 | 3.34E+07 | 3.39E+07 | 2.76E+07 | 3.21E+07 | 3.31E+07 | 3.54E+07 | 2.51E+07 | 2.01E+07 | 2.71E+07 | 4.89E+07 | 4.59E+07 | 3.11E+07 | 3.56E+07 | 3.84E+07 | 4.39E+07 | 2.63E+07 | 3.28E+07 | 3.47E+07 |
| Glutamine | normalized to 13C labeled internal control | 1.84E+00 | 1.20E+00 | 2.23E+00 | 3.45E+00 | 3.59E+00 | 2.83E+00 | 2.53E+00 | 1.50E+00 | 1.35E+00 | 6.88E-01 | 1.45E+00 | 1.11E+00 | 2.47E+00 | 4.79E+00 | 1.42E+01 | 3.91E+00 | 3.91E+00 | 2.89E+00 |
| Glucose-6-phosphate | normalized to 13C labeled internal control | 2.44E+00 | 5.81E+00 | 5.18E+00 | 7.42E-01 | 7.35E-01 | 6.49E-01 | 4.78E-01 | 4.09E-01 | 4.33E-01 | 5.04E+00 | 2.54E+00 | 2.29E+00 | 3.04E-01 | 3.90E-01 | 5.27E-01 | 3.25E-01 | 2.58E-01 | 3.80E-01 |
| GGPP | normalized to OD600 | 3.47E+06 | 3.60E+06 | 4.00E+06 | 4.37E+06 | 5.72E+06 | 5.11E+06 | 3.91E+06 | 4.77E+06 | 3.94E+06 | 3.57E+06 | 5.19E+06 | 3.92E+06 | 7.03E+06 | 5.90E+06 | 4.18E+06 | 3.23E+06 | 5.47E+06 | 4.51E+06 |
| Fumarate | normalized to OD600 | 6.74E+07 | 4.47E+07 | 4.49E+07 | 1.16E+10 | 1.07E+10 | 1.33E+10 | 1.23E+09 | 1.02E+09 | 1.29E+09 | 4.29E+07 | 6.01E+07 | 5.43E+07 | 1.16E+10 | 1.13E+10 | 1.12E+10 | 1.11E+09 | 1.27E+09 | 1.19E+09 |
| Fructose 1_6-bisphosphate | normalized to OD600 | 2.38E+06 | 6.91E+07 | 9.36E+07 | 4.54E+05 | 7.63E+05 | 1.07E+06 | 1.86E+05 | 7.56E+04 | 3.02E+04 | 6.96E+07 | 1.14E+08 | 6.24E+07 | 3.54E+06 | 5.16E+05 | 0.00E+00 | 3.03E+05 | 1.03E+05 | 2.22E+06 |
| FAD | normalized to OD600 | 4.18E+06 | 1.47E+06 | 1.19E+06 | 1.90E+06 | 3.50E+06 | 3.64E+06 | 1.26E+06 | 0.00E+00 | 0.00E+00 | 1.58E+06 | 3.14E+06 | 2.65E+06 | 4.21E+06 | 6.36E+05 | 1.42E+06 | 2.77E+06 | 1.65E+06 | 1.69E+06 |
| dTTP | normalized to OD600 | 1.44E+06 | 8.70E+06 | 7.41E+06 | 6.02E+06 | 8.16E+06 | 8.02E+06 | 4.80E+06 | 1.54E+06 | 3.46E+06 | 1.37E+07 | 1.46E+07 | 8.66E+06 | 7.34E+06 | 3.07E+06 | 4.75E+06 | 9.18E+06 | 8.16E+06 | 9.92E+06 |
| Dihydroxyacetone phosphate | normalized to 13C labeled internal control | 2.36E+00 | 3.71E+00 | 3.94E+00 | 2.52E+00 | 2.60E+00 | 2.13E+00 | 1.84E+00 | 1.71E+00 | 1.86E+00 | 4.31E+00 | 3.30E+00 | 1.61E+00 | 1.67E+00 | 2.12E+00 | 3.20E+00 | 2.23E+00 | 2.00E+00 | 1.94E+00 |
| Deoxyribose-5-phosphate | normalized to 13C labeled internal control | 4.11E+00 | 4.71E+00 | 5.02E+00 | 2.81E+00 | 4.02E+00 | 3.51E+00 | 2.67E+00 | 4.39E+00 | 4.54E+00 | 3.24E+00 | 1.65E+00 | 1.92E+00 | 3.45E+00 | 2.34E+00 | 2.50E+00 | 4.29E+00 | 3.47E+00 | 2.39E+00 |
| dCTP | normalized to OD600 | 1.06E+06 | 4.13E+06 | 3.31E+06 | 2.65E+06 | 3.05E+06 | 3.22E+06 | 1.13E+06 | 6.21E+04 | 8.03E+04 | 5.42E+06 | 8.87E+06 | 3.15E+06 | 5.97E+06 | 4.09E+06 | 1.09E+05 | 3.14E+06 | 1.20E+06 | 1.48E+06 |
| CTP | normalized to OD600 | 1.49E+05 | 3.12E+06 | 5.11E+06 | 4.58E+06 | 1.84E+07 | 1.41E+07 | 4.02E+06 | 6.38E+05 | 2.66E+06 | 4.43E+06 | 1.01E+07 | 3.63E+06 | 1.53E+07 | 4.85E+06 | 5.65E+06 | 1.17E+07 | 4.56E+06 | 8.36E+06 |
| cMEPP | normalized to 13C labeled internal control | 4.16E+00 | 8.10E+00 | 7.97E+00 | 4.86E+00 | 4.88E+00 | 4.09E+00 | 5.27E+00 | 4.15E+00 | 4.84E+00 | 8.08E+00 | 4.09E+00 | 2.98E+00 | 3.06E+00 | 2.56E+00 | 1.60E+01 | 4.40E+00 | 3.72E+00 | 2.49E+00 |
| CDP-MEP | normalized to OD600 | 2.29E+05 | 1.26E+06 | 1.36E+06 | 1.68E+06 | 1.76E+06 | 4.09E+05 | 6.29E+04 | 6.32E+04 | 6.80E+04 | 1.80E+06 | 4.06E+05 | 3.61E+05 | 4.52E+05 | 5.53E+04 | 1.59E+06 | 5.30E+04 | 2.87E+05 | 4.97E+04 |
| Asparagine | normalized to OD600 | 6.32E+07 | 4.69E+07 | 4.69E+07 | 8.14E+07 | 7.87E+07 | 8.96E+07 | 9.63E+07 | 7.79E+07 | 7.88E+07 | 4.62E+07 | 5.81E+07 | 5.86E+07 | 9.55E+07 | 1.02E+08 | 8.45E+07 | 6.54E+07 | 8.24E+07 | 7.66E+07 |
| AMP | normalized to OD600 | 4.79E+06 | 3.79E+06 | 2.01E+06 | 0.00E+00 | 0.00E+00 | 0.00E+00 | 6.96E+06 | 3.28E+06 | 6.11E+06 | 1.50E+06 | 5.40E+06 | 1.58E+07 | 6.47E+05 | 0.00E+00 | 3.13E+05 | 5.36E+06 | 3.80E+06 | 2.60E+06 |
| Acetyl phosphate | normalized to OD600 | 4.36E+06 | 1.09E+07 | 1.48E+07 | 1.06E+07 | 7.76E+06 | 8.15E+06 | 7.69E+06 | 5.62E+06 | 4.25E+06 | 1.90E+07 | 1.05E+07 | 5.30E+06 | 8.15E+06 | 1.13E+07 | 7.81E+06 | 5.22E+06 | 2.65E+06 | 4.17E+06 |
| 3-Dehydroshikimate | normalized to OD600 | 2.56E+00 | 6.23E+00 | 6.48E+00 | 1.23E+01 | 1.10E+01 | 1.22E+01 | 5.55E+00 | 5.59E+00 | 7.28E+00 | 8.93E+00 | 2.67E+00 | 1.76E+00 | 6.88E+00 | 7.96E+00 | 7.38E+00 | 6.95E+00 | 6.18E+00 | 6.27E+00 |
| 3-Dehydroquinate | normalized to OD600 | 2.39E+06 | 4.11E+06 | 7.78E+06 | 1.30E+07 | 1.46E+07 | 1.14E+07 | 5.22E+06 | 6.63E+06 | 6.68E+06 | 1.58E+07 | 3.13E+06 | 1.20E+06 | 9.68E+06 | 1.25E+07 | 1.60E+05 | 4.31E+06 | 4.30E+06 | 3.37E+06 |
| 2_3-Dihydrobenzoic acid | normalized to OD600 | 3.20E+08 | 1.44E+09 | 1.06E+09 | 1.15E+08 | 1.27E+08 | 8.16E+07 | 8.66E+06 | 3.14E+06 | 1.73E+07 | 2.93E+09 | 6.92E+08 | 1.10E+08 | 4.00E+07 | 0.00E+00 | 1.19E+08 | 1.55E+07 | 1.15E+07 | 8.30E+06 |

Supplementary Table 3: Absolute concentration of selected metabolites in wild type and *pykΔectd* cells (Unit: mM), measured by spiking in known concentrations of isotopologues

| | Glucose | | Malate | | Pyruvate | |
|---|---|---|---|---|---|---|
| | Wild type | *pykΔectd* | Wild type | *pykΔectd* | Wild type | *pykΔectd* |
| PEP | 0.11±0.0040* | 0.10±0.010 | 3.6±0.29 | 1.2±0.16 | 3.3±0.35 | 0.75±0.040 |
| R5P | 0.68±0.0079 | 0.73±0.038 | 0.30±0.016 | 0.12±0.024 | 0.23±0.070 | 0.05±0.020 |
| ATP | 3.1±0.17 | 3.2±0.042 | 4.4±0.56 | 4.8±0.63 | 4.1±0.55 | 3.6±0.32 |
| ADP | 0.66±0.066 | 0.78±0.027 | 1.4±0.035 | 1.3±0.032 | 0.95±0.20 | 0.45±0.063 |
| AMP | 0.12±0.025 | 0.12±0.025 | ND[2] | ND[2] | 0.20±0.12 | 0.15±0.17 |

* Data represents mean ± standard error of the mean of n=3 replicates.

[1] Cells were grown at 37 °C liquid culture with vigorous shaking, in minimal defined media with the indicated carbon source at 1% (w/v) concentration (see Materials and Methods: Metabolomic analysis by LC-MS).

[2] AMP signals were too low to be determined under these conditions.
